# Sponging of glutamate at the outer plasma membrane surface reveals roles for glutamate in development

**DOI:** 10.1101/2020.03.24.005223

**Authors:** Vanessa Castro-Rodríguez, Thomas J. Kleist, Nicoline M. Gappel, Fatiha Atanjaoui, Sakiko Okumoto, Mackenzie Machado, Tom Denyer, Marja C. P. Timmermans, Wolf B. Frommer, Michael M. Wudick

**Affiliations:** Institute for Molecular Physiology, Heinrich Heine Universität Düsseldorf, Germany; Department of Soil and Crop Science, Texas A&M, College Station, TX, USA; Dep. Plant Biology, Carnegie Institution for Science, Stanford, CA, USA; Center for Plant Molecular Biology, University of Tübingen, Auf der Morgenstelle 32, 72076 Tübingen, Germany; Institute of Transformative Bio-Molecules (WPI-ITbM), Nagoya University, Chikusa, Nagoya 464-8601, Japan

**Keywords:** *Arabidopsis thaliana*, glutamate, root meristem, fluorescence, biosensor

## Abstract

Plants use electrical and chemical signals for systemic communication. Herbivory, for instance, appears to trigger local apoplasmic glutamate accumulation, systemic electrical signals and calcium waves that travel to report insect damage to neighboring leaves and initiate defense. To monitor extra- and intracellular glutamate concentrations in plants, we generated Arabidopsis lines expressing genetically encoded fluorescent glutamate sensors. In contrast to cytosolically localized sensors, extracellularly displayed variants inhibited plant growth and proper development. Phenotypic analyses of high-affinity display sensor lines revealed that root meristem development, particularly the quiescent center (QC), number of lateral roots, vegetative growth and floral architecture were impacted. Notably, the severity of the phenotypes was positively correlated with the affinity of the display sensors, intimating that their ability to sequester glutamate at the surface of the plasma membrane was responsible for the defects. Root growth defects were suppressed by supplementing culture media with low levels of glutamate. Together, the data indicate that sequestration of glutamate at the cell surface either disrupts the supply of glutamate to meristematic cells and/or impairs localized glutamatergic signaling important for developmental processes.

**SIGNIFICANCE STATEMENT:** Glutamate is an important signaling molecule in animals and plants. The affinity-dependent effects of surface-displayed glutamate sensors on plant development intimate that glutamate levels in the vicinity of the plasma membrane play important roles in signaling in plants.

## INTRODUCTION

As a building block of proteins and central player in metabolism, glutamate is an essential amino acid for all known forms of life. In addition, many organisms also utilize L-glutamate as a signaling molecule. Glutamatergic signaling has been most intensively studied in the context of the animal central nervous system, where it serves as an excitatory neurotransmitter through activation of metabotropic and ionotropic glutamate receptors. In plants, glutamate is essential for the assimilation of nitrogen and for transamination (Forde and Lea, 2007). Low amounts of externally supplied glutamate are known to affect root architecture by impairing primary root growth while favoring the development of secondary roots (Walch-Liu *et al.*, 2006). Various lines of evidence implicate glutamate itself as a signaling molecule also in plants. For instance, mutants with lesions in *GLUTAMATE RECEPTOR-LIKE* (*GLR*) ion channel genes, encoding proteins for which glutamate can act as an agonist, show altered wound responses as a consequence of impaired propagation of wound-induced variation potentials (Mousavi *et al.*, 2013; Nguyen *et al.*, 2018). Notably, it has been shown that external application of glutamate to leaves elicits long-distance, calcium-based plant defense responses (Toyota *et al.*, 2018).

Genetically encoded biosensors that rely on changes in Förster resonance energy transfer (FRET) efficiency of two fluorescent proteins (FPs) have been engineered and used to dynamically report levels of several metabolites and signaling molecules in living cells (Frommer *et al.*, 2009; Okumoto *et al.*, 2012). By sandwiching a bacterial periplasmic binding protein (YbeJ) between enhanced cyan FP (ECFP) and Venus, a yellow FP (YFP) variant (Okumoto *et al.*, 2005), we engineered the first FRET-based fluorescent indicator proteins for glutamate (FLIPE). Subsequent systematic modification of first-generation glutamate sensors yielded a series of optimized sensors with varying ligand affinities and dynamic ranges (Deuschle *et al.*, 2005). However, the optimized variants could not be used in neuronal cells since they did not traffic efficiently to the cell surface (Okumoto unpublished). Subsequently, the intensiometric single-fluorophore iGluSnFRs were engineered by insertion of a circularly permutated green FP (cpGFP) into YbeJ (Marvin *et al.*, 2013). Toyota *et al.*, (2018) successfully targeted iGluSnFR to the apoplasmic space to estimate glutamate concentrations after wounding of Arabidopsis (*Arabidopsis thaliana*) leaves (Toyota *et al.*, 2018). Because cpGFP-based sensors are particularly sensitive to pH changes, the use of ratiometric FRET-based sensors is advantageous for *in vivo* imaging (Barnett *et al.*, 2017). We therefore sought to deploy a suite of ratiometric FLIPE variants spanning a wide range of affinities to enable monitoring extra- and intracellular levels of glutamate in Arabidopsis.

Plants expressing cytosolic versions of FLIPE showed wild type-like growth phenotypes. Unexpectedly, when FLIPEs were anchored in the plasma membrane and displayed extracellularly, transformants were severely stunted and showed defects in root and flower development, as well as growth. The severity of the phenotype depended on the affinity of the sensors, and symptoms could be ameliorated by supplementation with glutamate, consistent with a model wherein surface-displayed sensors interfere with glutamate availability close to the plasma membrane, thereby disrupting growth and development.

## RESULTS

### Expression of extracellularly displayed FLIPEs causes stunting and lethality

With the intent of generating plants for quantitative imaging of glutamate dynamics, a suite of FLIPE affinity variants with dissociation constants ranging from nanomolar to millimolar concentrations of glutamate was introduced into Arabidopsis and either targeted to the cytosol (FLIPE^cyt^) or tethered to the extracellular face of the plasma membrane, which we refer to here as ‘display’ sensors (FLIPE^display^). FLIPE^display^ and FLIPE^cyt^ cassettes were constructed by sandwiching the *E. coli* periplasmic glutamate binding protein YbeJ between two spectral variants of the green fluorescent protein. Cell surface display was achieved by fusion of an endoplasmic reticulum (ER) targeting sequence to the N-terminus of the chimera and the mammalian platelet-derived growth factor (PDGF) receptor transmembrane spanning domain to its C-terminus (Figure S1). The sensors were expressed under control of the ubiquitous CaMV 35S promoter in *rdr6-11* gene silencing mutants (Deuschle *et al.*, 2006) and visually selected for fluorescence (Figure S2a). For lines expressing the extracellularly displayed FLIPE, we noticed a positive correlation between sensor dissociation constant (K_*d*_) and severe growth defects (Figure 1a). Among soil-grown plants, rosette sizes of plants expressing extracellularly displayed sensors with affinities of 600 nM – 100 μM were significantly smaller, with rosette areas 20 – 50 % of the *rdr6-11* controls (Figure 1b). Moreover, plants expressing FLIPE-100μ^display^, FLIPE-1m^display^ were the only transgenic lines that generated seeds, albeit with reduced frequencies for display versions (9-32%) relative to controls (99%) (Table 1). To test whether noxiousness depends on the affinity of the binding protein, new senors with intermediate affinities (~40 μM: FLIPE-40μ; ~250 μM: FLIPE-250μ, ~500 μM: FLIPE-500μ) were engineered by site directed mutagenesis, and lines that display these sensors at the cell surface were generated and phenotyped (FLIPE-40μ^display^, FLIPE-250μ^display^, and FLIPE-500μ^display^; Figure S1, Figure S3). The majority of the plants expressing display sensors with intermediate affinities was dwarfed. Plants expressing the wild-type YbeJ cassette (FLIPE-600n^display^) were most affected (95%; Figure 1c). Intermediate affinity mutants showed “intermediate” (up to 34%) or wild type (up to 15%) growth phenotypes (Figure 1c). The defects could either be caused by effects of the binding cassette at the surface of the plasma membrane, scavenging glutamate in the apoplasmic space, or by processes in the secretory system. To evaluate if secretory system effects play a role, or if surface display, rather than mere apoplasmic localization of the sensors, was a prerequisite for the defects, a secreted version of FLIPE-600n carrying only the ER targeting sequence but lacking the PDGFR transmembrane domain was engineered (FLIPE-600n^apo^, Figure S1). Different from the displayed FLIPE-600n version, the majority of FLIPE-600n^apo^-expressing plants showed wild type-like growth (87%), similar to plants expressing the cytosolic FLIPE version of the same affinity (FLIPE-600n^cyt^, Figure 1c, Figure S4a, b). Since both display and apoplasmic sensors have to pass through the secretory system, the noxious effects are likely due to extracellular glutamate scavenging. Moreover, the presence of the sensor in the apoplasmic space did not seem to cause defects, whereas tethering to the extracellular face of the plasma membrane was likely causative for the observed developmental and growth defects. Specifically, we hypothesize that display of high-affinity glutamate-binding FLIPEs at the surface of the plasma membrane affects glutamate levels in the vicinity of receptors or transporters that are important for proper growth and development.

**Figure 1.**
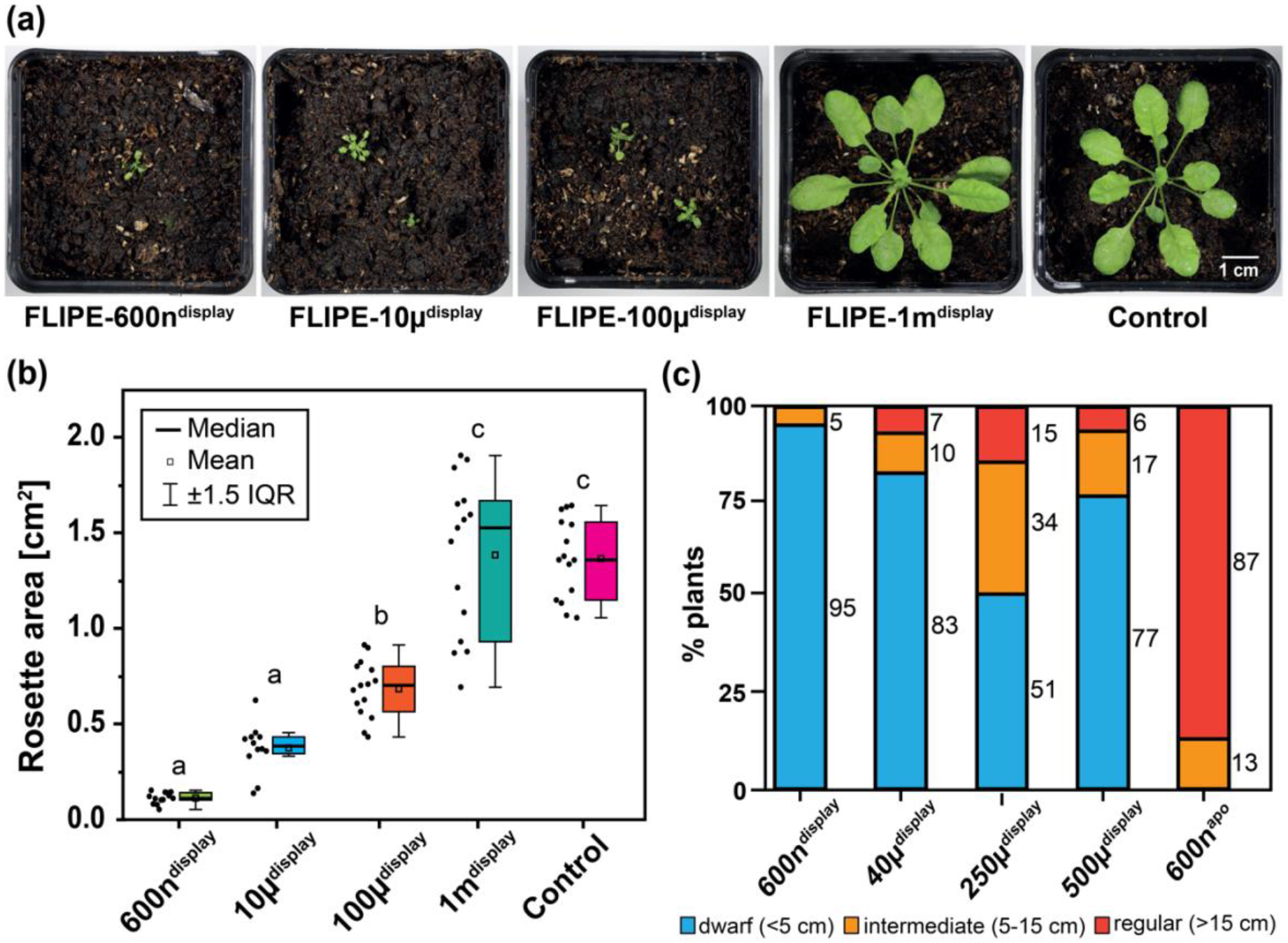
Phenotypic characterization of FLIPE sensor-expressing plants. (a) Representative phenotypes of 6-week-old plants grown on soil under short-day conditions, expressing FLIPE sensor affinity variants and the corresponding *rdr6-11* control. (b) Quantification of rosette areas of 3-week-old soil-grown plants expressing different FLIPE sensors (n ≥ 12 plants per genotype from 4 independent biological replicates). Compared to *rdr6-11* control plants, FLIPE-600n^display^, FLIPE-10μ^display^ and FLIPE-100μ^display^ plants showed significantly smaller rosettes, respectively, while the rosette sizes of cytosolic sensor variants and of the low-affinity FLIPE-1m^display^ line were not significantly different from the control (Tukey test, *P* < 0.001, letters indicate if samples are statistically indifferent (same letters) or different (different letters) from one another). (c) Phenotypic characterization into ‘dwarf’, ‘intermediate’ and ‘regular’ growth of several FLIPE^display^ and a FLIPE^apo^ version (n ≥ 21).

**Table 1.** Effect of FLIPE sensor variants on the lethality in Arabidopsis grown under long day conditions.

| sensor | # of plants transferred to soil | # of plants that survived after |  |  | % survival<sup>§</sup> | % lethality<sup>§</sup> |
| --- | --- | --- | --- | --- | --- | --- |
|  |  | week 2 | week 4 | week 6 |  |  |
| FLIPE-600n<sup>display</sup> | 55 | 12 | 3 | 0 | 0 | 100 |
| FLIPE-10μ<sup>display</sup> | 51 | 15 | 7 | 0 | 0 | 100 |
| FLIPE-100μ<sup>display</sup> | 65 | 19 | 11 | 6 | 9 | 90 |
| FLIPE-1m<sup>display</sup> | 73 | 34 | 30 | 23 | 32 | 68 |
| FLII<sup>81</sup>PE-1μ<sup>cyt</sup> | 107 | 106 | 106 | 106 | 99 | 1 |
| FLII<sup>81</sup>PE -10μ<sup>cyt</sup> | 109 | 108 | 108 | 108 | 99 | 1 |
| control (*rdr6-11*) | 103 | 102 | 102 | 102 | 99 | 1 |

<sup>§</sup> developmental stage: 6 weeks after germination, flowering and forming seeds

No obvious differences in fluorescence intensities were observed among plants expressing the full range of displayed FLIPE affinity variants (Figure S2a), indicating that sensor noxiousness was not caused by differences in expression levels. To exclude possible effects of the position of the fusion of the fluorescent proteins on the noxiousness of the sensors, we generated and analyzed plants expressing high affinity FLIPE sensors with altered sensor cassette topology, where ECFP was inserted into the YbeJ cassette at amino acid position 81 (FLII^81^PE-1μ (Deuschle *et al.*, 2005), FLII^81^PE-10μ, Figure S1). While plants expressing the display version of the FLII^81^PE-1μ sensor phenocopied plants producing the high affinity FLIPE sensors (*K*_*d*_ ~600 nM - 100 μM), plants expressing cytosolically targeted versions of the FLII^81^PE sensors (*K*_*d*_ ~1μM, 10 μM), were statistically indistinguishable from controls (Figure S4c, Figure S5, Table 1). To exclude the possibility that noxiousness of high affinity display sensors was caused by the membrane anchor, we replaced the PDGFR transmembrane domain with a polypeptide based on a previously described membrane anchor (Martinière *et al.*, 2018), yielding FLIPE-600n^display^-TM26 (Figure S1). Stable expression of the TM26-anchored FLIPE resulted in severely dwarfed plants, that were indistinguishable from plants expressing PDGFR-anchored sensors of the same affinity (Figure S4d), providing further evidence for the hypothesis that membrane display is imperative for noxiousness.

To determine the subcellular localization of the FLIPEs carrying different targeting signals, spinning disk confocal microscopy was performed on roots of FLIPE-expressing plants. FLIPE sensors lacking targeting signals expectedly localized to the cytosol (Figure 2a, b, Figure S6). While the localization of the display sensors was mostly limited to the plasma membrane (Figure 2d-f, Figure S6), we occasionally also observed apparent ER localization. Localization in ER-like structures was also observed for the apoplasmic sensor as would be expected for heterologous proteins entering the secretory pathway. However, although FLIPE-600n^apo^ was present in internal membranes (likely representing the ER), noxiousness was not observed, excluding a role along the secretory pathway (Figure 1c, Figure S4e).

**Figure 2.**
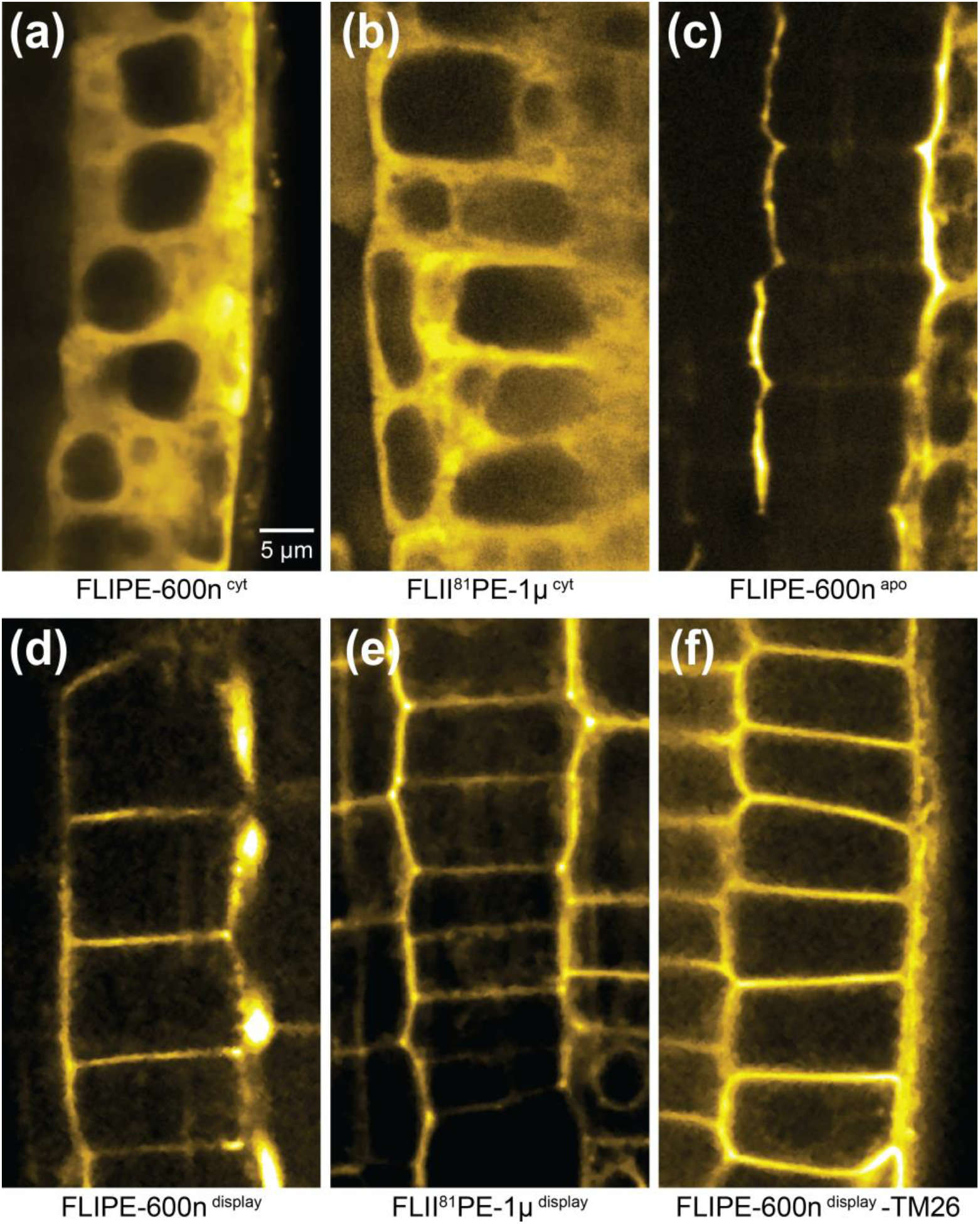
Subcellular localization of FLIPE sensors in the root proximal elongation zone. Representative confocal images of 3-5-day-old plants expressing FLIPE-600n^cyt^ (a), FLII^81^PE-1μ^cyt^ (b), FLIPE-600n^apo^ (c), FLIPE-600n^display^ (d), FLII^81^PE-1μ^display^ (e) or FLIPE-600n^display^-TM26 sensors (f). While sensors lacking additional targeting sequences accumulate throughout the cytosol (a, b), the apoplasmically-expressed sensor showed an accumulation in the cell-wall space and ER-like endomembranes (c), and the display sensors were predominantly localized to the plasma membrane but could also be detected in ER-like structures (d-f). Excitation and exposure times were independently adjusted to allow qualitative comparisons.

Surface display was evaluated in two ways: (*i*) by quantitative analysis of the sensor response, and by (*ii*) testing protease sensitivity. Consistent with the predicted surface display of the sensors, concentration-dependent changes in the fluorescence ratio were observed in response to perfusion of roots from FLIPE-10μ^display^ expressing plants with a *K*_*d*_ of ~10 μM (similar to the *K*_*d*_ of the sensor; Figure S7). Protease sensitivity assays provided additional evidence for the surface display of FLIPE-1m^display^. In contrast to the stable fluorescence from the cytosolic FLII^81^PE-1μ^cyt^ sensor, which was unaffected by preincubation with the protein synthesis inhibitor (cycloheximide, CHX), followed by proteinase K digestion in tobacco leaf protoplasts, fluorescence of FLIPE-1m^display^-expressing protoplasts decreased by approximately one third relative to protoplast treated with CHX alone (t = 42 min, Figure S8). These two sets of experiments support the display sensor topology wherein the ligand-binding domain of YbeJ faces the apoplasmic space and therefore is exposed to exogenously supplied protease.

Noxiousness might also be explained by differential proteolytic processing of cytosolic or apoplasmic sensors by endogenous proteases, i.e., by favoring cleavage of cytosolic variants. Indeed, in addition to polypeptides corresponding to the full-length versions of sensors targeted to the cytosol, plasma membrane or apoplasm, protein gel blots also revealed fragments of lower apparent molecular masses migrating in similar patterns, albeit for all sensors regardless of their subcellular localization (Figure S9). We therefore infer those proteolytic events most likely cannot explain the lack of noxiousness in plants expressing the high affinity sensor either in the cytosol or the apoplasm.

### Developmental defects in FLIPE^display^ lines

Phenotypic analyses of FLIPE^display^-expressing plants were performed to determine if the physiological effects of the FLIPE^display^ sensors impacted other aspects of plant development. Floral morphology was markedly different between FLIPE^display^ and control plants or cytosolic versions. Typically, flowers from dwarfed FLIPE^display^ plants were characterized by deformed or missing petals and failed to produce seeds (Figure 3). Morphological phenotypes were also observed and characterized in roots. Specifically, the root length of FLIPE^display^ lines showed an inverse proportionality to the affinity of the sensors for glutamate (*i.e*., plants expressing sensors higher affinities for glutamate had shorter primary roots). Since the extracellular glutamate concentration impacts mitotic activity in the root apical meristem (RAM, (Walch-Liu *et al.*, 2006)), RAMs from select FLIPE lines were examined. The majority of roots stably expressing the high-affinity FLIPE-600n^display^ sensor were severely deformed, likely due to aberrant growth and irregular cell divisions (Figure S10). Consequently, we were unable to unambiguously identify cell types and tissue layers in >80% of FLIPE-600n^display^-expressing roots and therefore excluded them from further analyses. While plants expressing FLIPE-10μ^display^ and FLIPE-100μ^display^ sensors also showed defective cell division patterns in the root meristem that affected the quiescent center (QC), columella and root cap cells, none of the roots were as severely affected as the roots expressing the FLIPE-600n^display^ sensor (Figure 4, Figure S10). Five to six days after germination, the root tips of control seedlings typically consisted of five tiers of cells, one or two of which represent stem cell tiers while the remaining tiers of differentiated cells are characterized by an accumulation of starch granules (Kiss *et al.*, 1996). We observed similar cellular patterns in RAMs of plants expressing FLII^81^PE-1μ^cyt^, FLII^81^PE-10μ^cyt^, and FLIPE-1m^display^ sensors as in *rdr6-11* controls (Figure 4, Figure S5e). Although roots from plants expressing FLIPE-10μ^display^ and FLIPE-100μ^display^ showed disorganized QC and columella cells, more than 50% of the mutants displayed at least one stem cell tier (Figure 4). Phenotypes of FLIPE^display^ plants are consistent with defects in the stem cell niche (affecting the overall number of cells), causing reduced root growth. As the above-ground tissues of plants expressing high-affinity FLIPE^display^ variants were severely dwarfed, it is possible that high-affinity FLIPE^display^ sensors also impact the development of the shoot meristem. Though not studied in detail here, together with the observed alterations in flower morphology, the findings tentatively point to a role of glutamate in developmental processes in meristems.

**Figure 3.**
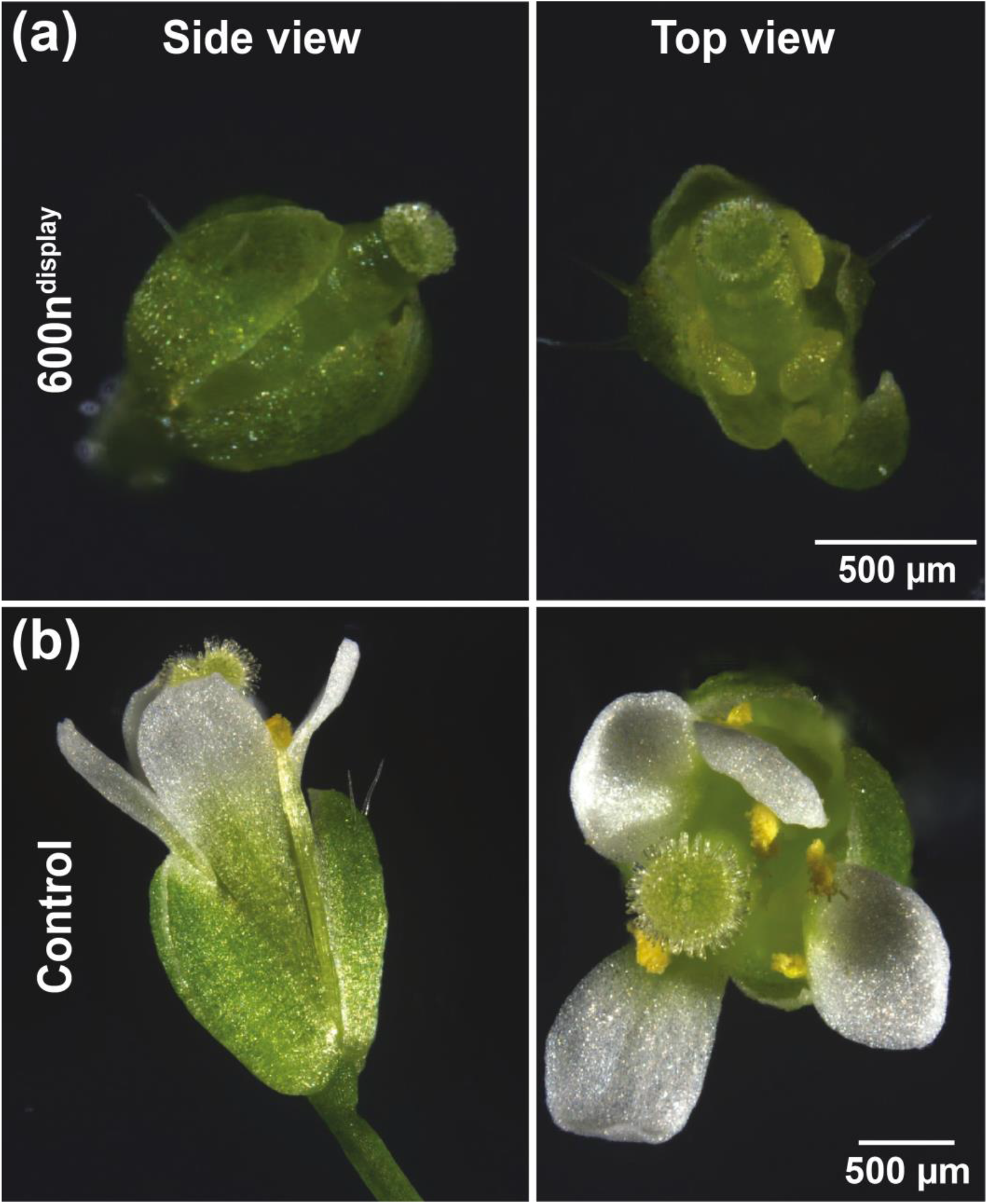
Phenotype of flowers from Arabidopsis control plants and lines expressing FLIPE-600n^display^. Representative flowers from (a) FLIPE-600n^display^ or (b) control (*rdr6-11*) plants.

**Figure 4.**
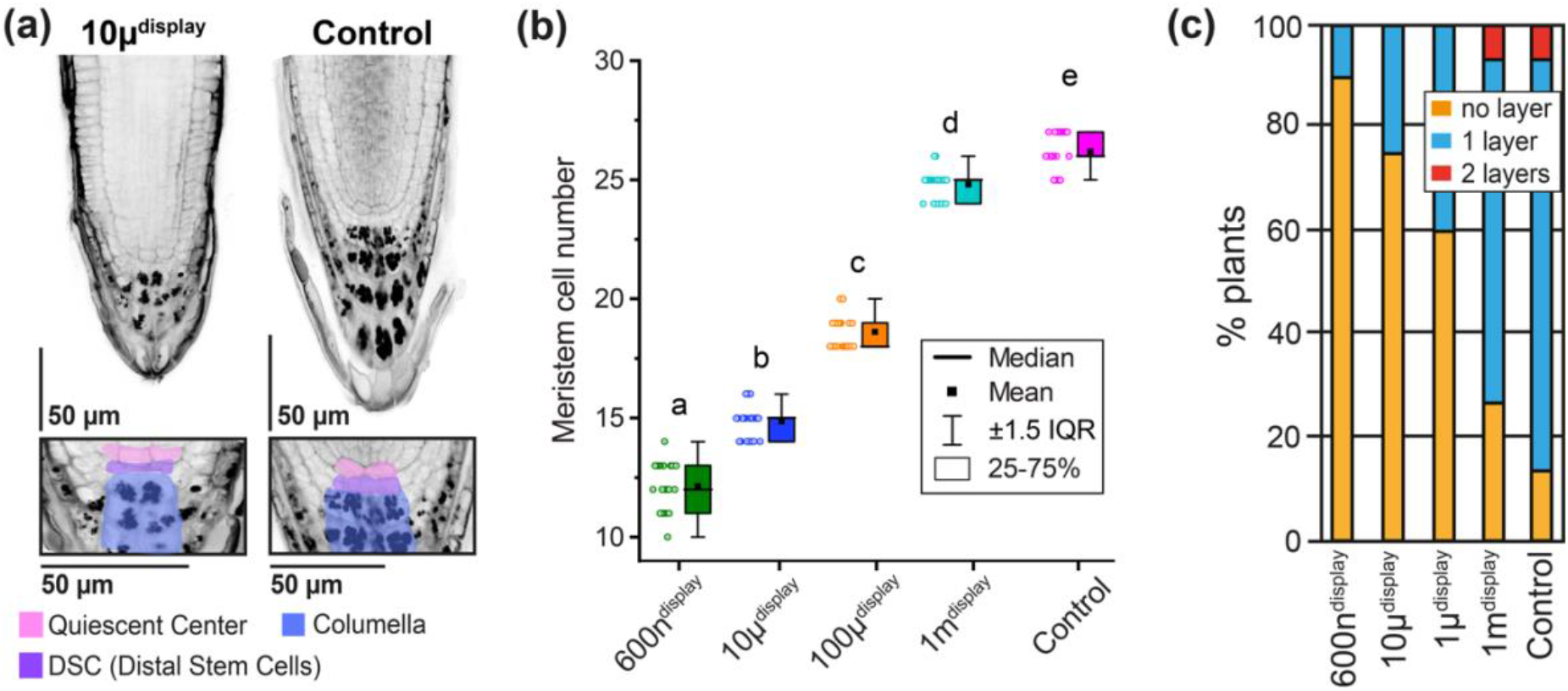
Effect of FLIPE sensor expression on the root apical meristem and the distal stem cell niche. (a) Representative root tips of plants expressing display and cytosolic FLIPE variants grown on AM media for 3-5 days after germination (top panel) and zoom-in of respective proximal meristems (lower panel). The color code highlights the Quiescent Center (QC, pink), Distal Stem Cells (DSC, purple) and Columella cells (blue). (b) Comparison of the size of the meristematic zone of the different FLIPE sensor lines, displaying the number of meristematic cells between QC and root elongation zone. Meristem cell numbers were statistically different in FLIPE-600n^display^, FLIPE-10μ^display^, FLIPE-100μ^display^, compared to control (*rdr6-11*; Tukey test *P* < 0.001, letters indicate if samples are statistically indifferent (same letters) or different (different letters) from one another). Data was acquired from 6 independent biological replicates, n = 17-21. (c) Percentage of roots with no clearly identifiable or undifferentiated DSC layer (yellow bar), one cell layer (blue bar) or two cell layers (red bar); experiment performed 3 independent times with similar results; n = 22.

### Glutamate supplementation mitigates root growth defects in FLIPE^display^ lines

The affinity-dependent effects of FLIPE^display^ lines indicate that the observed phenotypes may be caused by competitive binding (sponging) of extracellular glutamate at the cell surface. We therefore hypothesized that the growth defects could be suppressed by supplementing media with glutamate. Six-day-old seedlings expressing FLIPE-600n^display^ and untransformed controls were transferred from AM media to AM supplemented with glutamate. Exogenously supplied L-glutamate (eGlu) impairs primary root growth and increases the number of secondary roots (Walch-Liu *et al.*, 2006). Accordingly, while low concentrations of eGlu (0-25 μM) had little to no observable effect on root length of *rdr6-11* control plants, higher eGlu concentrations (> 100 μM) led to shorter primary roots (Figure 5a). In contrast, root lengths of FLIPE-600n^display^-expressing plants grown on media with 25-1000 μM eGlu were roughly twice as long compared to controls grown without glutamate supplementation. Therefore, supplementation with eGlu appeared to suppress the dwarf phenotype of plants expressing FLIPE-600n^display^ sensors, at least partially. Even at eGlu concentrations that inhibited root growth in *rdr6-11* plants, FLIPE-600n^display^-expressing plants showed increased primary root lengths compared to roots of plants grown in the absence of eGlu (Figure 5a). While glutamate concentrations > 25 μM elicited a previously described increase in the number of lateral roots in *rdr6-11* control plants (Walch-Liu *et al.*, 2006), FLIPE-600n^display^ plants did not show altered lateral root numbers (Figure 5b). Overall, eGlu treatment oppositely affected *rdr6-11* control and FLIPE-600n^display^-expressing plants. Sponging of glutamate at the cell surface could cause the defects in the meristems of plants expressing high-affinity FLIPE^display^ constructs either by limiting the supply of glutamate to meristematic cells or by preventing effective glutamate signaling processes in these cells. To distinguish between these two hypotheses, we analyzed cell type-specific levels of mRNAs associated with glutamate metabolism.

**Figure 5.**
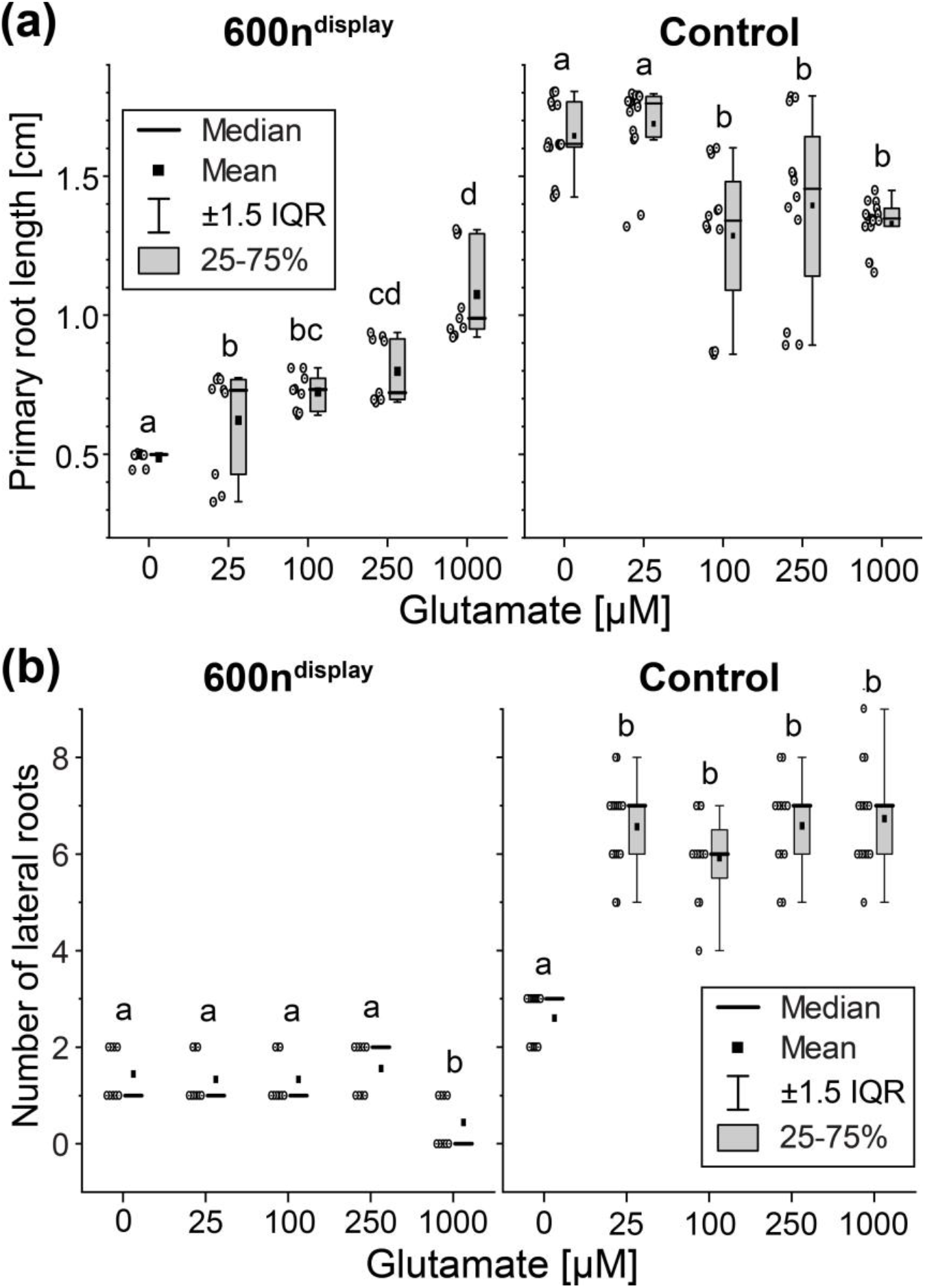
Supplementation with glutamate of FLIPE-600n^display^-expressing plants. (a) Primary root growth of vertically grown FLIPE-600n^display^ (left) or *rdr6-11* control (right) Arabidopsis seedlings 5 days after transfer to AM agar medium plates with increasing concentrations of glutamate. While roots of control plants did not significantly grow at [glu] ≥ 100 μM, roots lengths of FLIPE-600n^display^-expressing lines increased even at the highest [glu] tested (1 mM). (b) Number of lateral roots of FLIPE-600n^display^ (left) or *rdr6-11* control plants (right). Letters indicate if data were statistically indifferent (same letters) or different (different letters) from one another. Experiment was repeated three times independently with ≥ 3 roots per treatment yielding similar results.

### Transcripts of glutamate-associated genes are present in QC and surrounding cells

To profile of transcripts linked to glutamate-associated processes in the root tip meristem, single-cell RNA sequence (scRNA-seq) data from Arabidopsis root tips of 6-day-old seedlings clustered by cell type and developmental stage (Denyer *et al.*, 2019) were analyzed. Plants are able to synthesize glutamate through distinct enzymatic pathways, *i.e*., involving genes for *GLUTAMINE-α-OXOGLUTARATE AMINOTRANSFERASE* (*GOGAT*), *GLUTAMINE SYNTHETASE* (*GS*) and *GLUTAMATE DEHYDROGENASE* (*GDH*) in the presence of a nitrogen source (Forde and Lea, 2007). Our analysis revealed that transcripts of three different *GS* isoforms (*GS1;1*, *GS1;3* and *GS2*), as well as of *GOGAT2*, *GDH1* and the ammonium transporter *AMT1.1* were indeed detected in the QC and/or in surrounding cell layers (Figure S11), intimating that these cells are capable of synthesizing glutamate *de novo*. While transcripts of genes expected to be down-stream signaling targets of glutamate (*i.e.,* members of the *GLUTAMATE RECEPTOR-LIKE* (*GLR*) family of ion channels) showed low levels and lacked specific expression patterns (not shown), transcripts of a putative *LYSINE/HISTIDINE TRANSPORTER* gene (*LHT4*) were enriched in QC and surrounding cells (Figure S11). Homologs of LHT4 had previously been shown to mediate glutamate transport in rice (*Os*LHT1) and Arabidopsis (*At*LHT1) (Hirner *et al.*, 2006; Wang *et al.*, 2019). Taken together, transcriptomic data imply that root meristematic cells should be capable of synthesizing glutamate from inorganic ammonium, prompting us to hypothesize that aberrant glutamatergic signaling processes may underlie the observed noxiousness of FLIPE^display^ sensors. Together with the observations that only extracellularly displayed sensors caused developmental defects that could be partially alleviated by glutamate supplementation, we propose that sponging of eGlu directly at the outer face of the plasma membrane of meristematic cells has noxious effects on plant growth and development.

## DISCUSSION

With the initial intent to monitor glutamate dynamics *in planta*, Arabidopsis lines stably expressing a suite of Arabidopsis lines expressing fluorescent indicator proteins for glutamate (FLIPE) in the cytosol and displayed at the cell surface were generated. Surprisingly, plants expressing high-affinity FLIPE display variants were severely dwarfed and showed reduced or no fertility. The severity of the phenotype correlated positively with the affinity of the sensors for glutamate (*i.e*., high-affinity variants caused the most dramatic phenotypes). Prior characterization of the YbeJ binding domain revealed that it binds glutamate with an order of magnitude higher affinity than aspartate, and has substantially lower affinities for glutamine and asparagine (Okumoto *et al.*, 2005; Willis and Furlong, 1975). We therefore hypothesized that the phenotypes were caused by depletion of the glutamate pool at the cell surface due to the glutamate-binding activity of FLIPE constructs. Correspondingly, two high-affinity constructs (FLIPE-600n, FLII^81^PE-1μ) were only effective in causing developmental effects when anchored at the plasma membrane facing the apoplasm, while neither cytosolically nor apoplasmically targeted variants produced developmental or other phenotypic defects. Consistent with the hypothesis of glutamate depletion, supplementation of the media with extracellular glutamate partially restored growth in plants expressing extracellularly displayed, membrane-anchored high-affinity sensors. Sensor noxiousness could either be due to glutamate starvation in particular cell types, *i.e.,* in the root apical meristem, or perturbed glutamatergic signaling. Starvation is an unlikely cause, otherwise one may have expected that apoplasmic sponging would have the same noxious effect as found in display lines. The observation that transcripts for several glutamate-related genes are present in root meristematic cells, appears at odds with the hypothesis that glutamate starvation underpins FLIPE noxiousness. Moreover, sequestration of glutamate either in the cell wall space or the cytosol (in plants expressing sensors targeted to the apoplasm or cytosol, respectively) would be expected to interfere with metabolism in a qualitatively similar manner to FLIPE^display^ expression. Another possible explanation for the difference between cytosolic and display sensor effects could be that one of them is preferentially degraded, thus preventing (or causing) noxious effectors. Proteolytic degradation patterns of differently targeted FLIPEs were comparable and therefore most likely cannot explain (lack of) noxiousness. The preferred hypothesis is thus that FLIPE sensors interfere with glutamatergic signaling at the membrane surface, presumably by local buffering of glutamate pools in the vicinity of GLRs that may interfere with downstream signaling processes. Sequestration or buffering by transgenes has been described in plants, e.g., in the case of a genetically encoded cAMP-sponge (Sabetta *et al.*, 2019) or a parvalbumin-derived calcium buffer (Huang *et al.*, 2017).

For glutamate to function as a signaling molecule, it presumably must be kept at low resting levels in the extracellular space, particularly in the vicinity of glutamate receptors. In animals, this is well-established and achieved by the action of multiple sets of transporters in pre- and postsynaptic neurons, as well as adjacent glial cells (Mahmoud *et al.*, 2019). Interestingly, the effect of YbeJ-based glutamate indicators (iGluSnFR) on glutamate signaling in mice cortex astrocytes was recently studied. In a striking analogy to our findings, the authors come to the conclusion that extracellularly displayed sensors can act as glutamate buffers that reduce the extrasynaptic glutamate concentration, thereby slowing down glutamate uptake and yielding shorter synaptic transporter currents (Armbruster *et al.*, 2020).

A main difference between plants and animals, however, is that the volume of the synaptic cleft is extremely small, thus requiring only small amounts of glutamate to be released and subsequently taken up again for desensitization, while the apoplasmic space between plant cells is many orders of magnitudes larger, suggesting a need for different mechanisms. Therefore, efficient glutamate uptake and redistribution mechanisms are particularly important in plants, which use glutamate as a major transport form of nitrogen. Indeed, while the concentrations of most amino acids in plants may change substantially during the diurnal cycle, glutamate in the leaf apoplasm appears to be homeostatic between 0.3 – 1.3 mM, depending on the species (Forde and Lea, 2007). Glutamate receptor-like proteins (GLRs) have previously been implicated in meristem maintenance, developmental regulation and glutamatergic signaling processes in roots (Li *et al.*, 2006; Vincill *et al.*, 2013; Singh *et al.*, 2016; Singh and Chang, 2017). Similar to FLIPE^display^-expressing plants, rice plants carrying mutations in *GLR3.1* displayed short roots with reduced QC cell numbers and an impaired cell division activity of the RAM (Li *et al.*, 2006). Further evidence that glutamate-induced, GLR-mediated signaling is important for root development comes from the observation that glutamate receptor agonists negatively affected the primary root length and the number of secondary roots (Singh and Chang, 2017). The authors reported that, similar to the effects of glutamate resupply on FLIPE-expressing lines, exogenous application of glutamate was able to restore primary root growth as well as the number of lateral roots, further indicating that root development relies on glutamate signaling processes. A recent publication determined the apparent *in vitro* dissociation constant for glutamate of the GLR3.3 ligand binding domain with *K*_*d*_ = 2.2 ± 0.5 μM (Alfieri *et al.*, 2020). This value is very similar to the apparent affinities for glutamate of the FLIPE-600n and FLII^81^PE-1μ constructs we used (with *K*_*d*_ ≈ 600 nM and 1 μM, respectively), and might support the idea of competitive binding of the glutamate by the FLIPE constructs.

Future studies should examine if the buffering of glutamate occurs in specific membrane (nano-) domains, as membrane compartmentalization and the establishment of functional sub-membrane domains were shown to be critical for plant cell signaling events (Gronnier *et al.*, 2017; Chu *et al.*, 2021).

The noxious effects of FLIPE^display^ sensors render them unsuitable for glutamate imaging, for instance during wounding. Analysis of glutamate dynamics are thus best performed with untethered apoplasmic glutamate sensors with appropriate affinities. Our study cautions that targeting of YbeJ-derived sensors might impact plant physiology. If able to be deployed inducibly, extracellularly displayed YbeJ chimeras may however provide a novel tool to study glutamatergic signaling in plants.

In summary, our data suggest that the level of glutamate specifically at the cell surface is important for glutamatergic signaling, thus requiring tight control via transport processes. Moreover, our data may support a role of glutamate in the regulation of developmental processes, in particular in root, shoot and floral meristems.

## MATERIAL AND METHODS

### FLIPE constructs for plant transformation

FLIPE surface display constructs carrying tandemly fused murine Ig kappa-chain (IgK) signal peptides, the YbeJ coding sequence (also called GltI; NP_415188, amino acid residues 29 - 302), and the transmembrane domain of the platelet-derived growth factor receptor (PDGFR) were constructed as described previously (Okumoto *et al.*, 2005). Except for the FLII^81^PE constructs (see below), the YbeJ coding sequence was sandwiched between sequences encoding enhanced cyan fluorescent protein (ECFP) and Venus, a yellow FP. The resulting construct was excised using *BamH*I/*Xho*I, and subcloned into *BamH*I/*Sal*I sites of the plant expression vector CF203 (a pPZP212 derivative (Hajdukiewicz *et al.*, 1994) containing the CaMV 35S‐promoter, GFP5(S65T) gene and a rbcS terminator (kindly provided by C. Fankhauser, Lausanne)). Affinity mutants were designed from wild-type YbeJ (*K*_*d*_ ≈ 600 nM) carrying different substitutions in position 179: A179R (*K*_*d*_ ≈ 10 μM), A179V (*K*_*d*_ ≈ 100 μM), and A179W (*K*_*d*_ ≈ 1 mM) (Okumoto *et al.*, 2005) and additional intermediate affinities that were generated in this work: A179Q (*K*_*d*_ = 40 ±0,87 μM; mean ±s. e. m. *n* = 9), A179Y (*K*_*d*_ = 256 ±4,63 μM; mean ±s. e. m. *n* = 9) and A179N (*K*_*d*_ = 514 ±8,92 μM; mean ±s. e. m. *n* = 9). Fluorescent indicator proteins for glutamate (FLIPE) sensors were named FLIPE-x^*display*^ or FLIPE-x^*cyt*^ depending on their expression at the surface (display) or in the cytosol (cyt), respectively. The resulting plasmids were called FLIPE-600n^display^; FLIPE-10μ^display^; FLIPE-40μ^display^; FLIPE-100μ^display^; FLIPE-250μ^display^; FLIPE-500μ^display^; FLIPE-1m^display^ and FLIPE-600n^cyt^. The FLIPE-600n^display^-TM26 construct was generated using In-Fusion technology by replacing the PDGFR transmembrane domain with a previously described transmembrane domain (Martinière *et al.*, 2018). Additionally, a FLIPE-600n^display^-based construct lacking the C-terminal PDGFR transmembrane domain was generated (FLIPE-600n^apo^) for apoplasmic targeting. The cytosolic and display variants of the FLII^81^PE-1μ and FLII^81^PE-10μ sensor are based on a previously published construct (Deuschle *et al.*, 2005), where the ECFP is inserted into the YbeJ cassette after amino acid 81, which was subsequently subcloned into the CF203 vector. Generated plasmids with complete sequence information will be made available via Addgene upon publication.

*Agrobacterium tumefaciens* GV3101 was transformed with the binary vectors and *Arabidopsis thaliana rdr6-11* silencing mutants were transformed by floral dipping. Seeds were surface-sterilized with 70 % ethanol for 5 min followed by 30 % sodium hypochlorite for 10 min and rinsed with sterile, deionized water. Transformants were selected on AM media (half-strength Murashige and Skoog (MS) basal salt mixture including MES buffer (Duchefa Biochemies; cat No. M0254.0050) supplemented with 1 % plant agar (pH 5.7) (Sigma, cat. no. 9002-18-0) and 1 % sucrose. Seedlings were grown vertically under long-day light conditions (16 h light, 8 h dark) at a light intensity of about 100 μmol/m^2^ sec for 3 days. To identify plant expressing the sensor, seedlings were preliminarily screened for GFP fluorescence (bandpass excitation filter at λ_ex_ = 470/40 nm, bandpass emission filter at λ_em_ = 525/50 nm, with a beam splitter at λ_em_ = 495 nm) using a fluorescence stereo zoom microscope ZEISS Axio Zoom.V16, which can detect ECFP and Venus fluorescence. Plant transformation with FLIPE sensors was carried out independently at least three times and yielded comparable results.

### *In vitro* characterization of FLIPE affinity variants

Affinity mutants carrying the substitution A179Q, A179Y and A179N were created by site-directed mutagenesis, and cloned into pRSET vector for protein expression in *E. coli*, adding an N-terminal 6xHis-tag to the constructs. Sequences were verified by sequencing. pRSET-FLIPE constructs were transferred to *E. coli* BL21(DE3) Gold (Stratagene) by chemical transformation. Colonies were inoculated in LB-media containing carbenicillin (Sigma Aldrich; cat No. 4800-94-6), 0.2 % lactose (Merck; cat No. 7660.0250) and 0.05 % glucose (Sigma Aldrich; cat No. 14431-43-7) and expressed for 2 h at 37 °C, 220 rpm shaking and then 48 h at 20 °C in darkness. Cells were harvested by centrifugation for 30 minutes at 4 °C, 13,000 x *g.* Sediments were resuspended in 20 mM sodium phosphate buffer (pH 7.0) containing protease inhibitors and lysed by sonication [Branson Sonifier cell disruptor B15; 10 cycles x (15s ON/OFF); 60 % amplitude; 90 % duty cycle], after which the lysate was centrifuged for 1 h at 4 °C, 13,000 x *g*. Supernatants were purified by histidine-affinity-chromatography. 6xHis-tagged proteins were eluted from the beads by using imidazole (AppliChem; cat No. A1073,0500; UN3263). Ligand titration curves were performed by using a microplate reader TECAN™ Spark^®^. ECFP fluorescence was acquired at λ_ex_ = 430/10 nm and λ_em_ = 470/10 nm, respectively. Venus fluorescence was recorded with a bandpass excitation filter at λ_ex_ = 500/10 nm and a bandpass emission filter at λ_em_ = 530/10 nm. Intensity scans were performed at λ_ex_ = 430/10 nm and λem = 460-600/10 nm. All analyses were done using 20 mM sodium phosphate buffer, pH 7.0. The *K*_*d*_ of each FLIPE variant was determined by fitting to a single site binding isotherm as described previously (Okumoto *et al.*, 2005). Affinities were calculated from at least three independent protein extracts.

### Plant growth

Transgenic *Arabidopsis thaliana* (L.) Heynh. lines stably expressing display, cytosolic or apoplasmic FLIPE or FLII^81^PE constructs were grown in soil Floradur^®^ (Floragard Vertriebs-GmbH products, Oldenburg, Germany) in individual 2.5-inch pots. Pots were arrayed in a 4 × 8 grid in standard greenhouse flats (Hermann Meyer KG products, Germany) and transferred to growth chambers with short day (8 h light, 16 h dark) conditions at 21°C and 50-70 % relative humidity, unless stated differently. Plants were watered as needed, depending on growth stage. To determine rosette sizes, plants were imaged every week during 5 weeks until flowering. Size classification was carried out on plants grown under long day conditions. Images of the flowers from each transformant were acquired using a ZEISS Axio Zoom.V16 equipped with a ZEISS Axiocam 305 color camera. Image analysis to determine vegetative area and flower anatomy was performed using FIJI. Experiments were repeated four independent times, yielding comparable results.

### Subcellular localization of FLIPE sensors

Roots of 5–7-day-old seedlings expressing variously targeted FLIPE sensors were imaged using an Olympus IXplore SpinSR confocal microscope equipped with UPLSAPO 60x 1.3 NA silicone immersion oil objective (Olympus), 50 μm spinning disk, and Photometrics Prime BSI sCMOS camera. A 40 mW coherent OBIS LX 514 nm laser was used to excite YFP using a 445/514/640 nm dichroic and 534/23 nm emission filter for detection. Laser power and acquisition time were adjusted to accommodate differences in brightness among lines.

Additional experiments were performed to validate the localization of the FLIPE^display^ sensor facing the apoplasm.

Subcellular localization assays were adapted from (Martinière *et al.*, 2018) and repeated twice, yielding comparable results. Transient expression of the sensors was performed in leaves of 3–4-week-old *Nicotiana benthamiana* plants infiltrated with *Agrobacterium tumefaciens* strain GV3101 carrying constructs for FLIPE-1m^display^ and FLI^81^IPE-1μ^cyt^, as described below. In brief, 3 days past infiltration, protoplasts were obtained by digesting transformed leaves with 1.2% cellulose Onozuka R10 (Duchefa; cat No. C8001), 0.4 % macerozyme R10 (Duchefa; cat No. M8002) in 0.4 M mannitol and MES 20 mM (pH 6.5) for 6-8 h at room temperature in the dark by shaking at 1x *g*. Protoplasts were pelleted at 100 x *g* and rinsed with fresh buffer. Preincubation with 50 μM cycloheximide (CHX, Sigma Aldrich; cat No. 66-81-9) for 30 min was followed by digestion with proteinase K at 50 μg/mL and 50 μM CHX. Slides with protoplasts were imaged under a confocal microscope for at least 20 min. Venus fluorescence intensity was quantified using FIJI(Schindelin *et al.*, 2012).

### Glutamate supplementation of FLIPE seedlings

Glutamate supplementation experiments were performed by measuring the growth of seedlings transformed with FLIPE constructs on square petri dishes with AM agar medium supplemented with 1 % sucrose (Sigma Aldrich; cat No. 57-50-1) (as described above) covered with a layer of nylon mesh (Sefar Nitex 03-15/10). Three days after germination, seedlings on the mesh were transferred to an AM agar medium supplemented with 1 % sucrose and 0 μM, 25 μM, 100 μM, 250 μM or 1 mM L-glutamic acid monosodium salt monohydrate (Sigma Aldrich, cat No. 6106-04-3). Plants were analyzed in single blinded-experiments. After transfer, plates were rotated 90 ° and plant were grown for 7 days under long day conditions (16 h light, 8 h dark). Seedlings were photographed daily during glutamate treatment and roots were measured using ImageJ (rsb.info.nih.gov/ij/). Experiments were repeated three independent times and yielded comparable results.

### Root tip (mPS)-PI staining

Modified pseudo-Schiff propidium iodide (mPS-PI) staining was performed as described by (Truernit *et al.*, 2008), on root tips 5 days after germination. Samples were fixed in fixative solution [50 % methanol (Sigma Aldrich, cat No. 67-56-1) and 10 % acetic acid (Sigma Aldrich, cat No. 64-19-7)] at 4 °C overnight. Root tips were washed with water and incubated in 1 % periodic acid at room temperature for 40 min. Then, tissues were rinsed and incubated in Schiff’s reagent (Sigma Aldrich, cat No. 3952016) with propidium iodide (100 mM sodium metabisulphite and 0.15 N HCl; propidium iodide to a final concentration of 100 μg/mL freshly added) until samples were visibly stained. Roots were transferred to chloral hydrate solution [70 % chloral hydrate (Sigma Aldrich, cat No. 302-17-0), 10 % glycerol (Sigma Aldrich, cat No. 56-81-5)] and slides were prepared for microscope analysis adding drops of chloral hydrate solution and left covered overnight. The samples were examined with a 40x water immersion objective. Samples stained with propidium iodide were excited with a 561 nm argon laser with emission detection at 566-718 nm.

### Total protein extraction and protein gel blot analyses

Plant tissue was freshly harvested from 6–7-day-old *Arabidopsis thaliana rdr6-11* controls or plants expressing FLIPE-600n^apo^, FLII^81^PE-1μ^cyt^ or FLIPE-600n^dis^ constructs and ground to a fine powder in liquid nitrogen. The protein extraction was performed by adding 1 mL per g fresh weight ice-cold extraction buffer (50 mM Tris-HCl (pH 7.5), 150 mM NaCl, 10 % glycerol, 2 mM EDTA, 5 mM DTT, 1 % Triton X-100, protease inhibitor mix (Roche, 1 tablet)), followed by vigorous vortexing. The extracts were centrifuged (20000 x *g*, 30 min, 4 °C) and the supernatant was collected. A purified GFP-tagged protein served as control. Protein gel blotting procedures followed standard protocols. In brief, Laemmli buffer was added to the samples, which were denatured at 95 °C for 7 min. Samples were loaded and run on a 4 - 20 % precast polyacrylamide gel (BioRad, 4561094, 10 μL per sample, except FLIPE-600n^display^: 20 μL) and subsequently electroblotted onto a PVDF membrane for immunoanalysis. After transfer, membranes were blocked with 3 % milk powder and incubated with a GFP antibody from Abcam (catalog nr. ab13970, 1:5000 dilution, 4 h) or Roche (catalog nr. 11814460001, 1:1000 dilution, overnight). Both antibodies also detect the GFP variant Venus, which is part of the FLIPE sensor. Bound antibodies were visualized using the goat anti-chicken IgY (H+L) Alexa Fluor 488 (Thermo Scientific, catalog nr. A-11039, 1:5000 dilution, 2 h) according to manufacturer’s instruction and α-mouse-HRP (Sigma, catalog nr. A-9044, 1:50000 dilution, 2-3 h) as secondary antibodies, respectively. The fluorescence activity of the Alexa Fluor 488 was detected in tiling acquisition mode using the Zeiss Axio Zoom.V16 scope and excitation at 470/40 nm. For detecting the HRP secondary antibody, the ECL SuperSignal™ West Femto (ThermoScientific, 34094) was used with subsequent visualization on a LAS-4000 Mini bioimager (GE-Healthcare Life Sciences). Images were analyzed with ImageJ (rsb.info.nih.gov/ij/). Contrast and intensities were adjusted separately for the extracted samples and the control to avoid signal saturation in the control.

### Quantitative imaging

Quantitative imaging was carried out as described previously (Chaudhuri *et al.*, 2011). In brief, 7-day-old seedlings grown on hydroponic media were mounted on coverslips (24×50 mm No. 1½, VWR) using medical adhesive (Stock No. 7730, Hollister) to restrict movement (Deuschle *et al.*, 2006). Chambers used for screening were made with plastic clay (Sculpey, www.sculpey.com) and were variable in size and volume (1–2 mL). For qualitative analyses, the clay chamber was filled with hydroponic medium, perfusion tubing was mounted, and roots were perfused at 3 mL. min^−1^. Ratio imaging was performed on an inverted fluorescence microscope (DM IRE2, Leica) with a QuantEM digital camera (Roper) and a ×20 oil objective (HC PL APO ×20/0.7IMM CORR, Leica, Germany). Dual emission intensities were recorded simultaneously using a DualView with a dual CFP/YFP-ET filter set [high transmission modified Magnetron sputter-coated filter sets ET470/24m (470 indicates the emission wavelength, /24 indicates bandwidth); ET535/3, Chroma, USA] and Slidebook software (Intelligent Imaging Innovations Inc., USA]. Data was analyzed with ImageJ (rsb.info.nih.gov/ij/).

## ACKNOWLEDGEMENTS

We would like to thank Rüdiger Simon’s group for assistance with PI staining of roots and the center for advanced imaging (CAi) at HHU for help with image acquisition. This research was funded by grants from Deutsche Forschungsgemeinschaft (DFG, German Research Foundation) under Germany’s Excellence Strategy – EXC-2048/1 – project ID 390686111 and Deutsche Forschungsgemeinschaft (DFG, German Research Foundation) SFB 1208 – Project-ID 267205415 – as well as the Alexander von Humboldt Foundation to WBF.

## AUTHOR CONTRIBUTIONS

Conceptualization: VCR, MMW, SO, WBF, TK; Methodology: FA, NG, SO, VCR, TK, TD, MT, MMW; Investigation: FA, NG, SO, VCR, TK, MM, MMW; Writing: VCR, TK, WBF; MMW Supervision: WBF, TK, MMW.

The authors declare no conflict of interest.

## Notes

### Competing Interest Statement

The authors have declared no competing interest.

### Summary of Updates

We performed additional experiments, generated additional control constructs and lines and phenotyped them. We also noted an error in the previous description of two cytosolic FLIPE sensors (600n and 10u). They represented different versions of the sensor in which the eCFP is inserted into the backbone of the glutamate binding protein. This error has no impact on the conclusions. This was corrected here and some of the additional controls included cytosolic FLIPE-600n with terminal fusions, which are formally more suited as controls for the display versions. We added a control where we targeted the intramolecular fusion sensor FLI81IPE-1u to the cell surface. The resulting lines behaved similar as the ones carrying the terminal fusions.

