## Supplementary Figures for "Sponging of glutamate at the outer plasma membrane surface reveals roles for glutamate in development"

### SUPPLEMENTAL FIGURES

Vanessa Castro-Rodríguez[1], Thomas J. Kleist[1], Nicoline M. Gappel[1], Fatiha Atanjaoui[1], Sakiko Okumoto[2], Mackenzie Machado[3], Tom Denyer[4], Marja C. P. Timmermans[4], Wolf B. Frommer[1,5] & Michael M. Wudick[1,*]

Affiliations:

[1] Institute for Molecular Physiology, Heinrich Heine Universität Düsseldorf, Germany

[2] Department of Soil and Crop Science, Texas A&M, College Station, TX, USA

[3] Dep. Plant Biology, Carnegie Institution for Science, Stanford, CA, USA

[4] Center for Plant Molecular Biology, University of Tübingen, Auf der Morgenstelle 32, 72076 Tübingen, Germany

[5] Institute of Transformative Bio-Molecules (WPI-ITbM), Nagoya University, Chikusa, Nagoya 464-8601, Japan

#### SUPPLEMENTARY FIGURES

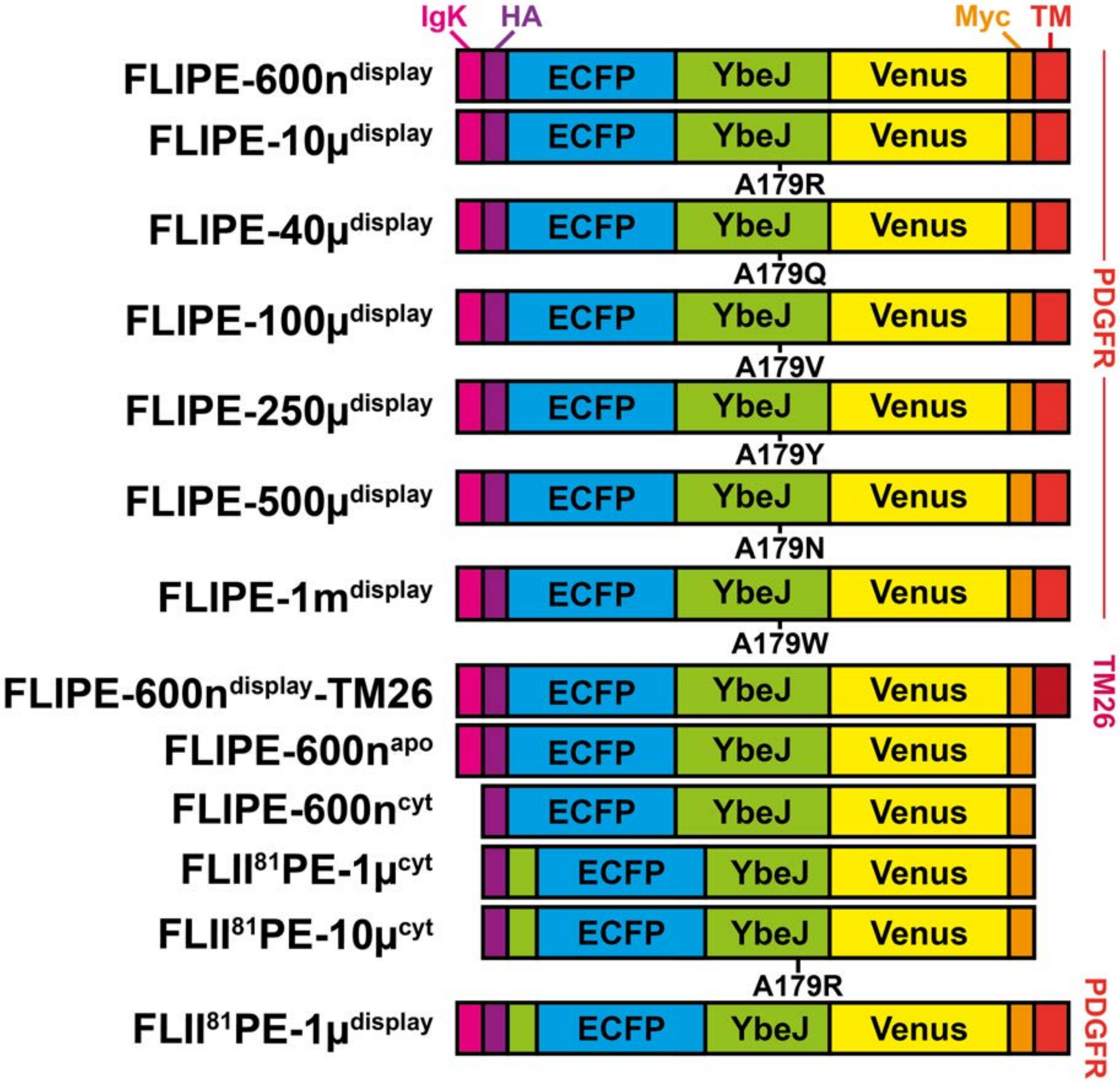


**Figure S1. Schematic representation of FLIPE and FLII$^{81}$PE sensor variants.** FLIPE$^{display}$ and FLIPE$^{cyt}$ cassette containing the enhanced cyan fluorescent protein (ECFP, blue box) and the yellow FP Venus (yellow box), flanking the periplasmic glutamate binding protein YbeJ from *E. coli*. (green box). Display sensor constructs contain sequences for an IgK ER targeting sequence (pink box), an HA tag at the 5'-end (purple box), and a c-Myc tag (orange box) followed by the transmembrane spanning domain (TM) of the human PDGF receptor (red box) or a 26 amino acid transmembrane domain (TM26, (Martinière *et al.*, 2018)). Amino acid changes in YbeJ at position 179 yielding different glutamate binding affinities are highlighted. The sequence of the 600n$^{apo}$ construct is identical to the 600n$^{display}$ construct but missing the PDGFR domain. FLII$^{81}$PE-1µ and FLII$^{81}$PE-10µ sensors were generated by inserting the ECFP downstream amino acid 81 into the YbeJ cassette, yielding a sensor with optimized dynamic range (Deuschle *et al.*, 2005). Sensor targeting: apo - apoplasmic, cyt - cytosolic, display - plasma membrane.

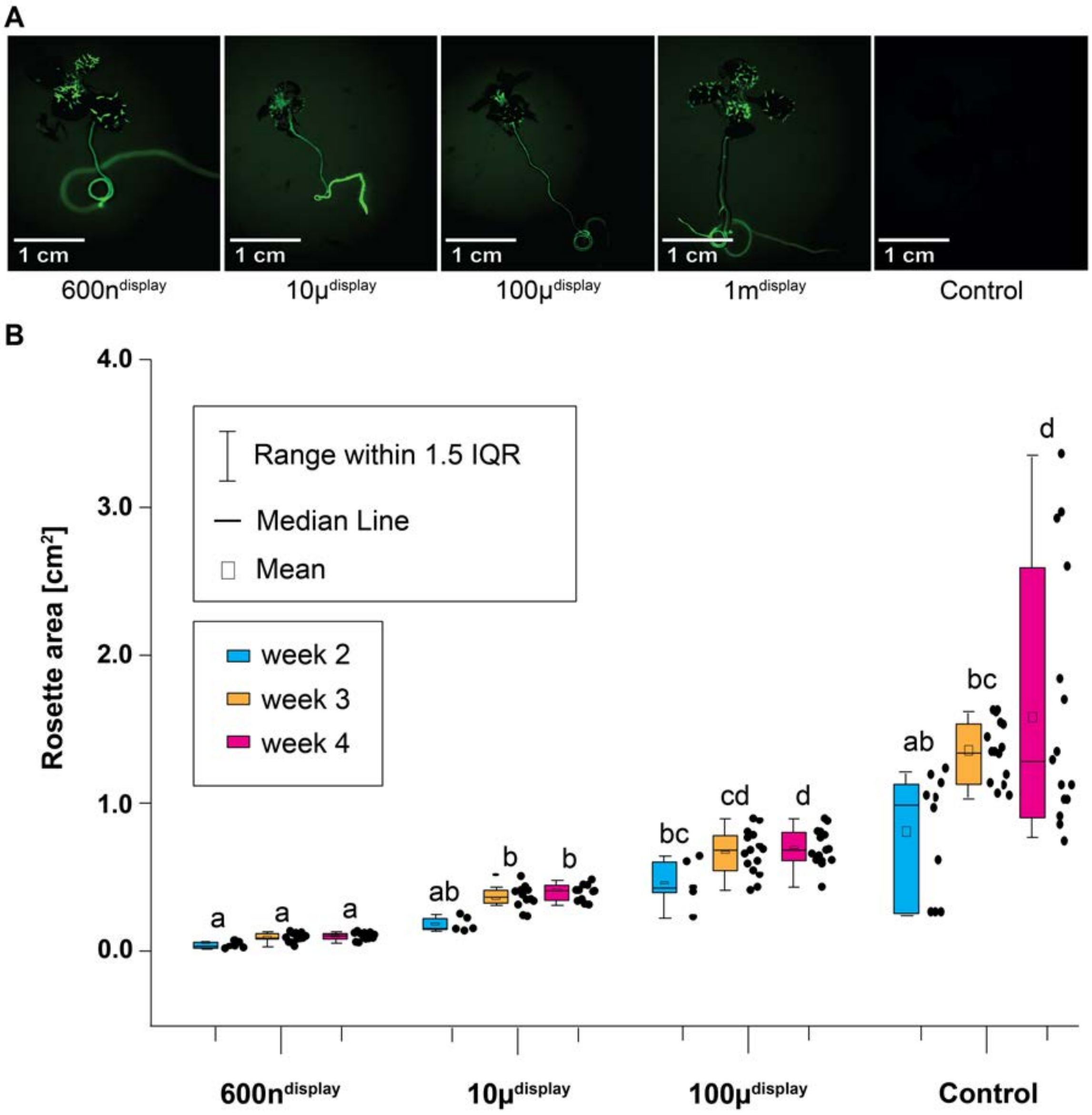


**Figure S2. Fluorescence in *rdr6-11* Arabidopsis seedlings expressing different FLIPE sensors and detailed growth characterization**. **(a)** 14-day-old Arabidopsis seedlings growing in half-strength MS media. Fluorescence of FLIPE$^{display}$, FLIPE$^{cyt}$ and non-transformed control plantlets (*rdr6-11*) was recorded using a ZEISS Axio Zoom.V16 stereo zoom microscope and GFP filter settings ($\lambda_{ex}$ = 470/40 nm and $\lambda_{em}$ = 525/50 nm, with a $\lambda$ = 495 nm beam splitter). All Arabidopsis experiments involving stably expressing FLIPE variants were carried out on $T_1$ plants derived from floral dip genetic transformation of $T_0$ plants. **(b)** Quantification of rosette areas of 2, 3 and 4-week-old soil-grown plants expressing different FLIPE sensors (n ≥ 12 plants per genotype from 4 independent biological replicates). In contrast to *rdr6-11* controls and plants expressing cytosolically targeted FLIPE sensors, plants expressing FLIPE-600n$^{display}$, FLIPE-10µ$^{display}$ and FLIPE-100µ$^{display}$ versions did not significantly increase rosette seize beyond week 3 (Tukey test, $P < 0.001$). Letters indicate if samples are statistically indistinguishable (same letters) or significantly different (different letters) from one another. The values for 3-weeks grown plants are the same as in the Figure 1.

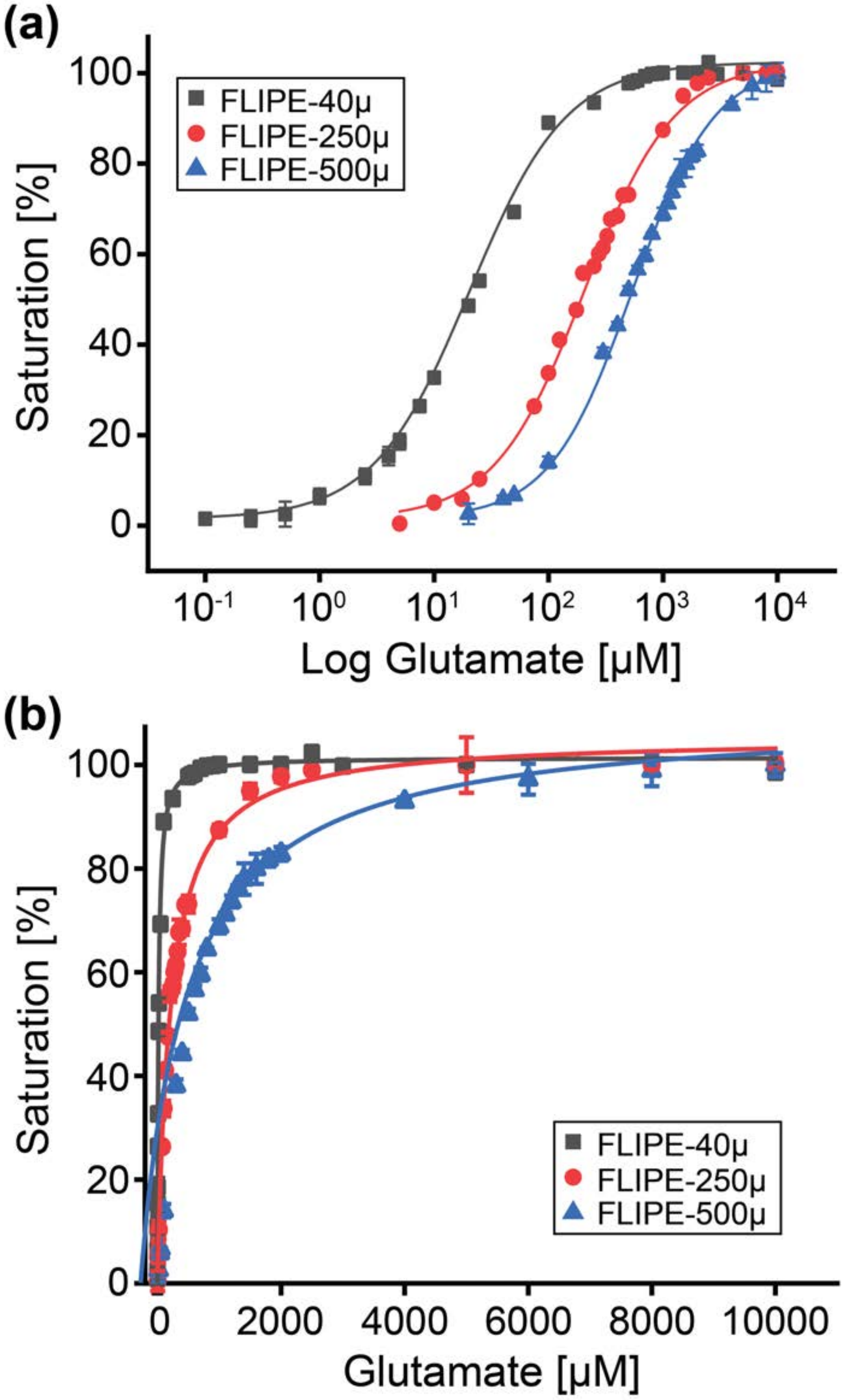


**Figure S3.** ***In vitro*** **saturation curves from FLIPE-40µ, FLIPE-250µ and FLIPE-500µ sensors.** Logarithmic **(a)** or linear **(b)** display of glutamate binding isotherms of affinity mutant FLIPE-40µ (black), FLIPE-250µ (red) and FLIPE-500µ (blue) sensors displayed as. Saturation of the sensors with different glutamate concentrations was performed by fluorimetry using proteins purified from *E. coli*. Glutamate binding was determined as previously described (Okumoto *et al.*, 2005). Data are presented as the average of three independent experiments.

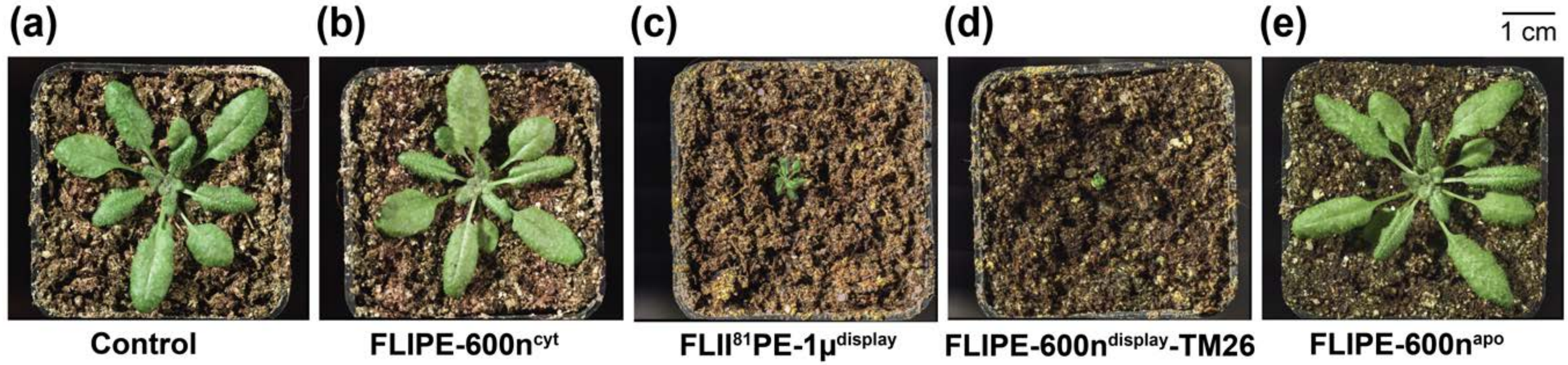


**Figure S4. Phenotypic characterization of additional FLIPE sensor-expressing plants. (a-e)** Representative phenotypes of 4–5-week-old plants grown on soil under prolonged short-day conditions (10 h light, 14 h dark at 21°C and 60 % relative humidity) expressing FLIPE sensor affinity variants in the cytosol (cyt), at the plasma membrane (display) or in the apoplasm (apo), compared to the corresponding *rdr6-11* control. White balance, exposure, and contrast were adjusted individually for images using Lightroom (Adobe) to improve visibility.

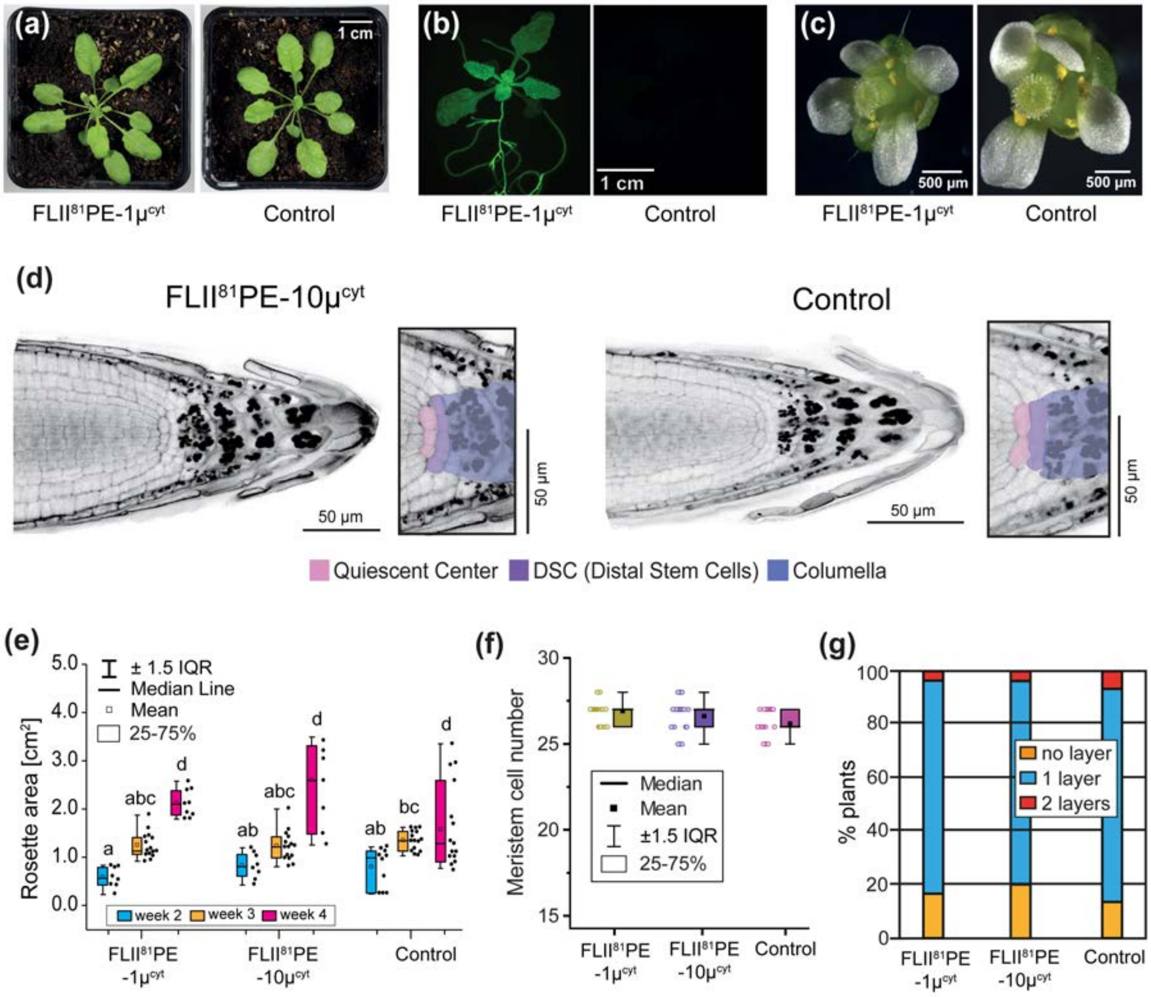


**Figure S5. Characterization of plants expressing different FLII$^{81}$PE variants. (a)** Representative phenotypes of 6-week-old plants grown on soil under short-day conditions, expressing the FLII$^{81}$PE-1μ$^{cyt}$ sensor and the corresponding *rdr6-11* control. **(b)** 14-days-old Arabidopsis seedlings growing in half-strength MS media. Fluorescence was recorded using a ZEISS Axio Zoom.V16 stereo zoom microscope and GFP filter ($\lambda_{ex}$ = 470/40 nm and $\lambda_{em}$ = 525/50 nm, with a beam splitter at $\lambda$ = 495 nm). **(c)** Representative flower from FLII$^{81}$PE-1μ$^{cyt}$-expressing plants in comparison to *rdr6-11* control. **(d)** Representative root tips of plants expressing FLII$^{81}$PE-10μ$^{cyt}$ (left panel) and control (*rdr6-11*, right panel) grown on half-strength MS medium for 3-5 days after germination and zoom-in of respective proximal meristems (rectangular panel). The color code highlights the Quiescent Center (QC, pink), Distal Stem Cells (DSC, purple) and Columella cells (blue). **(e)** Quantification of rosette areas of 2, 3 and 4-week-old soil-grown plants expressing FLII$^{81}$PE-1μ$^{cyt}$, FLII$^{81}$PE-10μ$^{cyt}$ or the *rdr6-11* control ($n \geq 12$ plants per genotype from 4 independent biological replicates) showed no statistically significant difference between the two genotypes. **(f)** Comparison of the size of the meristematic zone of the FLII$^{81}$PE-1μ$^{cyt}$ and FLII$^{81}$PE-10μ$^{cyt}$ sensor lines as compared to the *rdr6-11* control, displaying the number of meristematic cells between QC and root elongation zone. Data was acquired from 6 independent biological replicates (n = 17-21). **(g)** Percentage of FLII$^{81}$PE-1μ$^{cyt}$, FLII$^{81}$PE-10μ$^{cyt}$ or control (*rdr6-11*) roots with no clearly discernible DSC layer (yellow bar), one cell layer (blue bar) or two cell layers (red bar); experiment performed 3 independent times; n = 22. All images and values for control lines are reproduced from the corresponding main figures of the manuscript or from Figure S3.

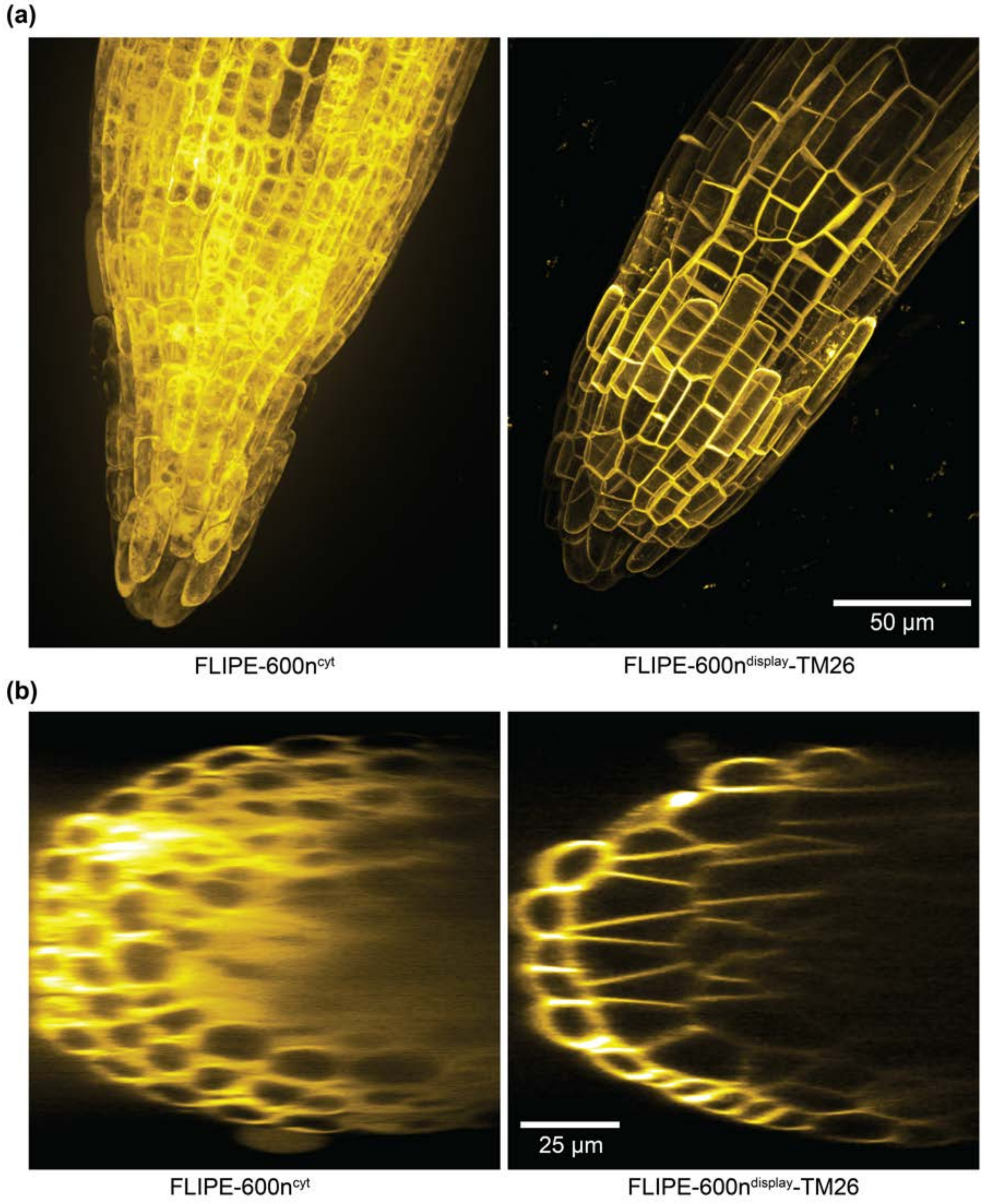


**Figure S6. Subcellular localization of FLIPE-600n targeting variants in roots. (a)** Maximum z-projections of roots from plants stably expressing FLIPE-600n$^{cyt}$ (left panel) or FLIPE-600n$^{display}$-TM26 (right panel) showing a clearly distinguishable distribution pattern for cytosolic and plasma membrane localization, respectively. **(b)** Orthogonal projections of the same confocal microscopy data. Acquisition settings were not identical due to differences in fluorescence intensities therefore are not quantitatively comparable. Excitation and emission settings were optimized for YFP fluorescence detection. Images are shown in ‘Yellow Hot’ lookup table (ImageJ).

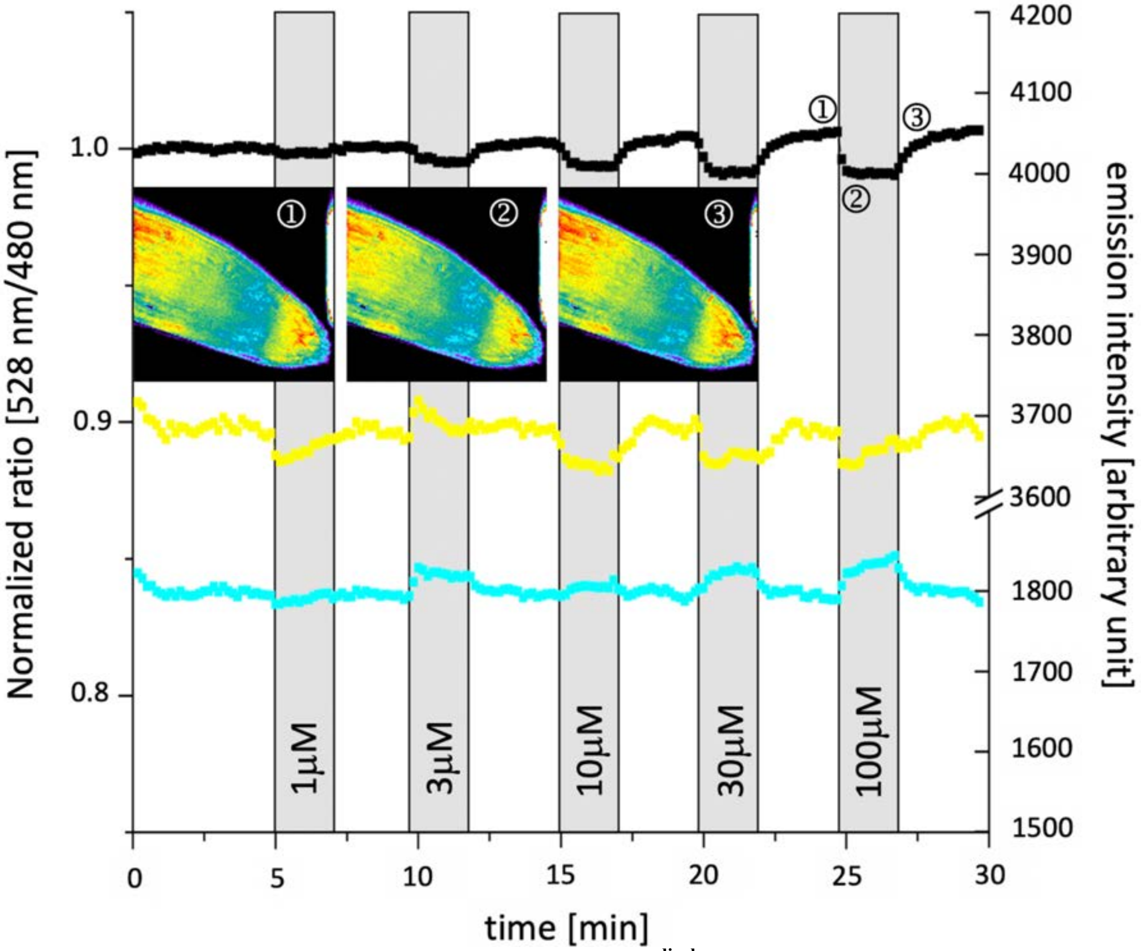


**Figure S7. *In vivo* characterization of FLIPE-10μ$^{display}$ in Arabidopsis roots.** Glutamate-induced FRET changes in intact roots of a 7-day-old plant. The FLIPE-10μ$^{display}$ FRET sensor response was recorded following the perfusion of a root from stably transformed *rdr6-11* plants with glutamate at concentrations of 1 μM, 3 μM, 10 μM, 30 μM and 100 μM, respectively, as indicated by the grey bars. The black trace represents the normalized ratio of Venus fluorescence intensity divided by ECFP fluorescence intensity. The individual recordings for Venus or ECFP fluorescence intensities are shown as yellow and blue traces, respectively. The inset depicts pseudo color FRET ratio images of the measured root tip at three indicated time points after stimulation with 100 μM glutamate. The apparent *in vivo* $K_d$ of 3-10 μM closely matches to the apparent *in vitro* affinity of the FLIPE-10μ$^{display}$ sensor. Of note, no response for cytosolic sensors could be detected, possibly due to insufficient rates of uptake of glutamate relative to its metabolism (Okumoto *et al.*, 2008).

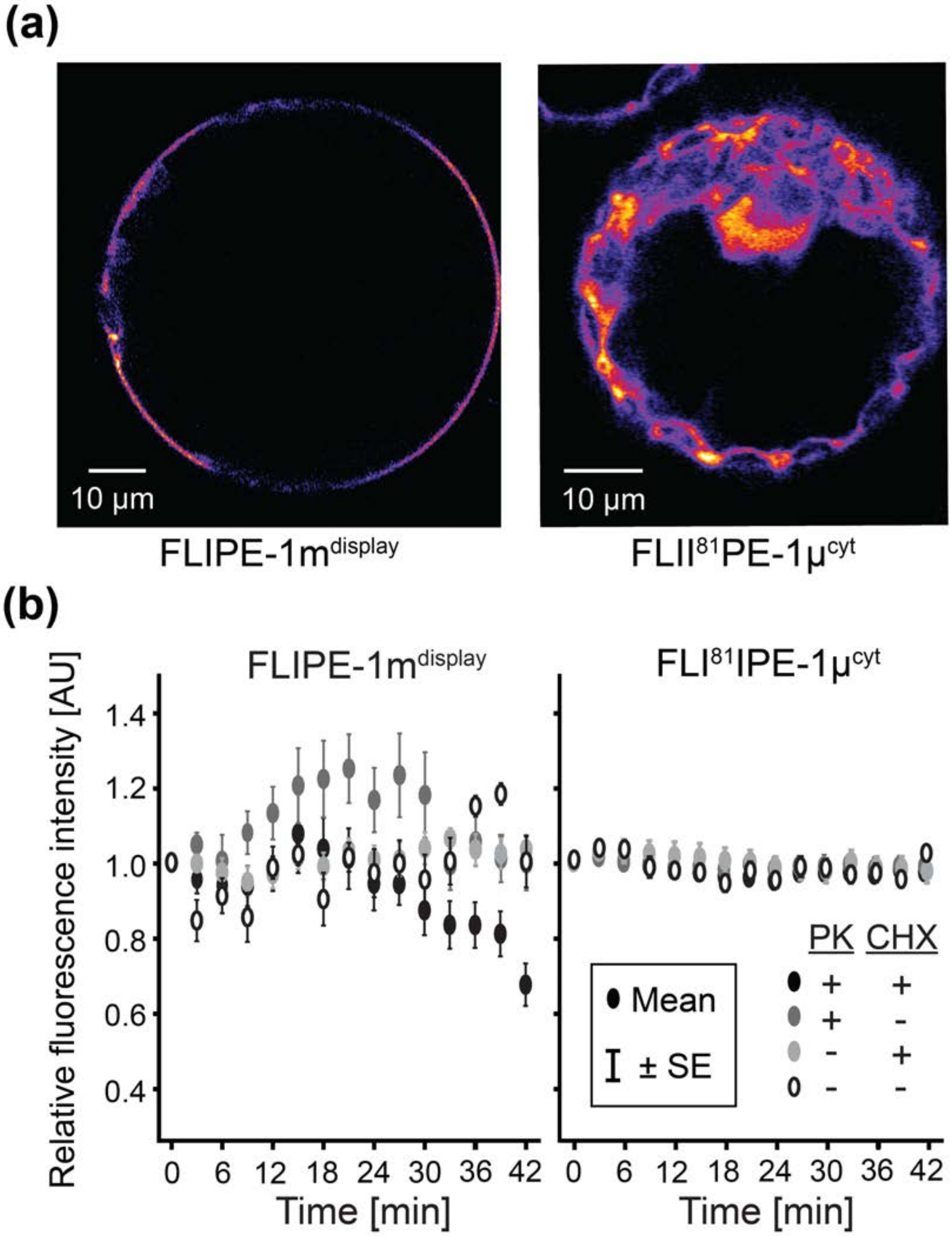


**Figure S8.** ***In vivo*** **proteinase K digestion assay of protoplasts expressing FLIPE sensors. (a)** Fluorescence from intact tobacco leaf protoplasts expressing FLIPE-1m$^{display}$ (left image) or FLII$^{81}$PE-1µ$^{cyt}$ (right image). Fluorescence of Venus is shown using a pseudo-color 'fire' lookup table. **(b)** Recording of protoplast fluorescence over time in the presence (+) or absence (-) of proteinase K (PK) and/or the protein synthesis inhibitor cycloheximide (CHX). In protoplasts treated with PK and CHX, a discernible decrease in Venus fluorescence was only observed for the plasma membrane-localized FLIPE-1m$^{display}$ sensor (left panel) but not for the cytosolically targeted FLII$^{81}$PE-1µ$^{cyt}$ (right panel), which is consistent with exposure of the FLIPE-1m$^{display}$ sensor to the apoplasm and its accessibility to proteolytic cleavage.

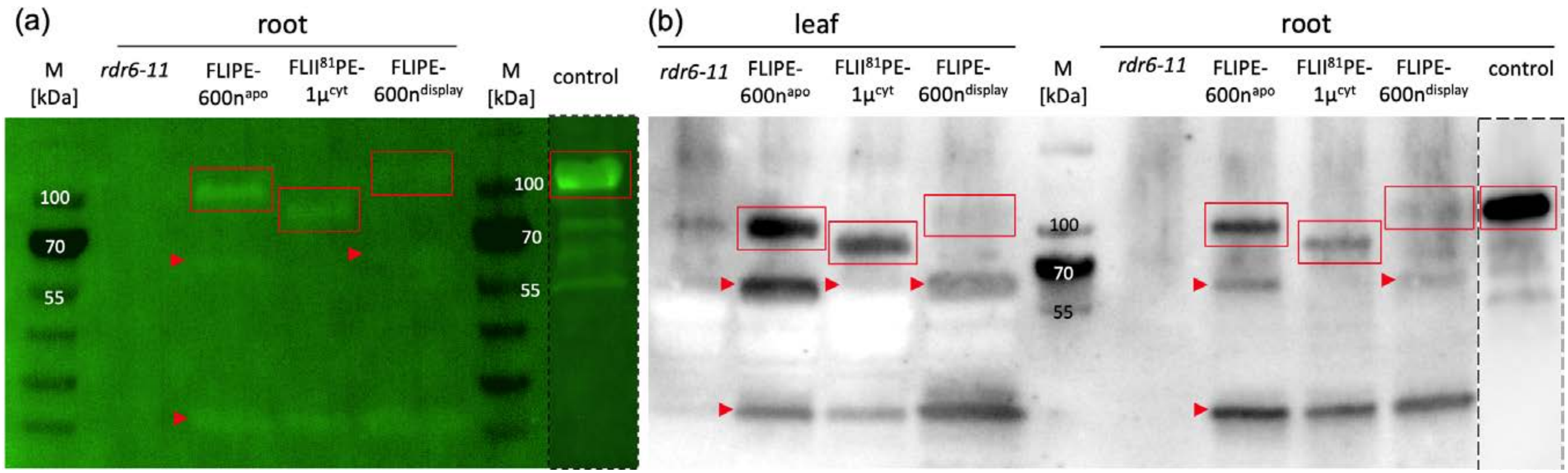


**Figure S9. Western blot analyses of different plant-expressed FLIPE variants. (a)** AlexaFluor 488 labeled Western blot of total proteins extracted from roots of *rdr6-11* control plants or plants expressing different sensor variants. **(b)** Chemiluminescence-based detection of total protein extracts from leaf and root tissue of *rdr6-11* control plants or plants expressing different sensor variants. Since the extraction protocol was not optimized for membrane proteins, intensities from full-length FLIPE-600n$^{dis}$-derived peptides appear expectedly weaker on the blots. Full-length polypeptides are highlighted by red rectangles. Additional polypeptides of lower molecular mass that likely resulted from proteolysis are indicated by red arrowheads. Calculated molecular masses: FLIPE-600n$^{apo}$: 91.1 kDa, FLII$^{81}$PE-1µ$^{cyto}$: 85 kDa, FLIPE-600n$^{display}$: 96.3 kDa, control: 103.3 kDa. To avoid saturation, contrast adjustment of images depicting the control were treated separately from the rest of the picture (indicated by dashed lines) M – protein standard marker, control – GFP-tagged soluble protein.

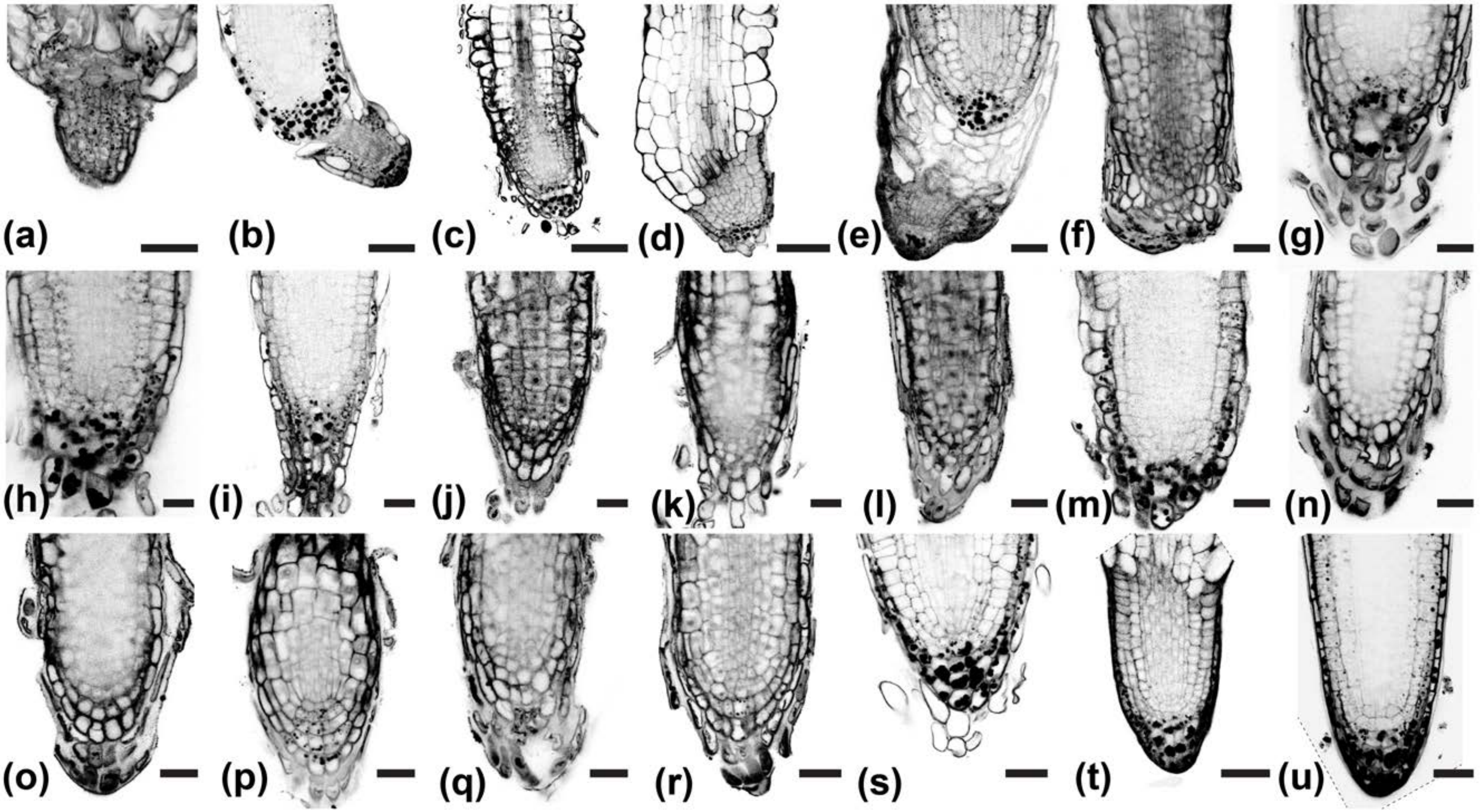


**Figure S10. Effect of FLIPE-600n$^{display}$ sensor expression on root development. (a-u)** Representative root tips of plants stably expressing the high-affinity display glutamate sensor, grown on half-strength MS medium and analyzed 3-5 days after germination. Sensor expression resulted in severely stunted and/or deformed roots in the majority of cases. In the most affected roots (a-d), the cellular identity and organization was too altered to reliably determine cell numbers or layers. Those roots were excluded from further analyses. We did not observe similarly severe phenotypes in roots expressing the other FLIPE variants (data not shown). Dotted lines indicate regions cropped due to image rotation. Scale bars = 20

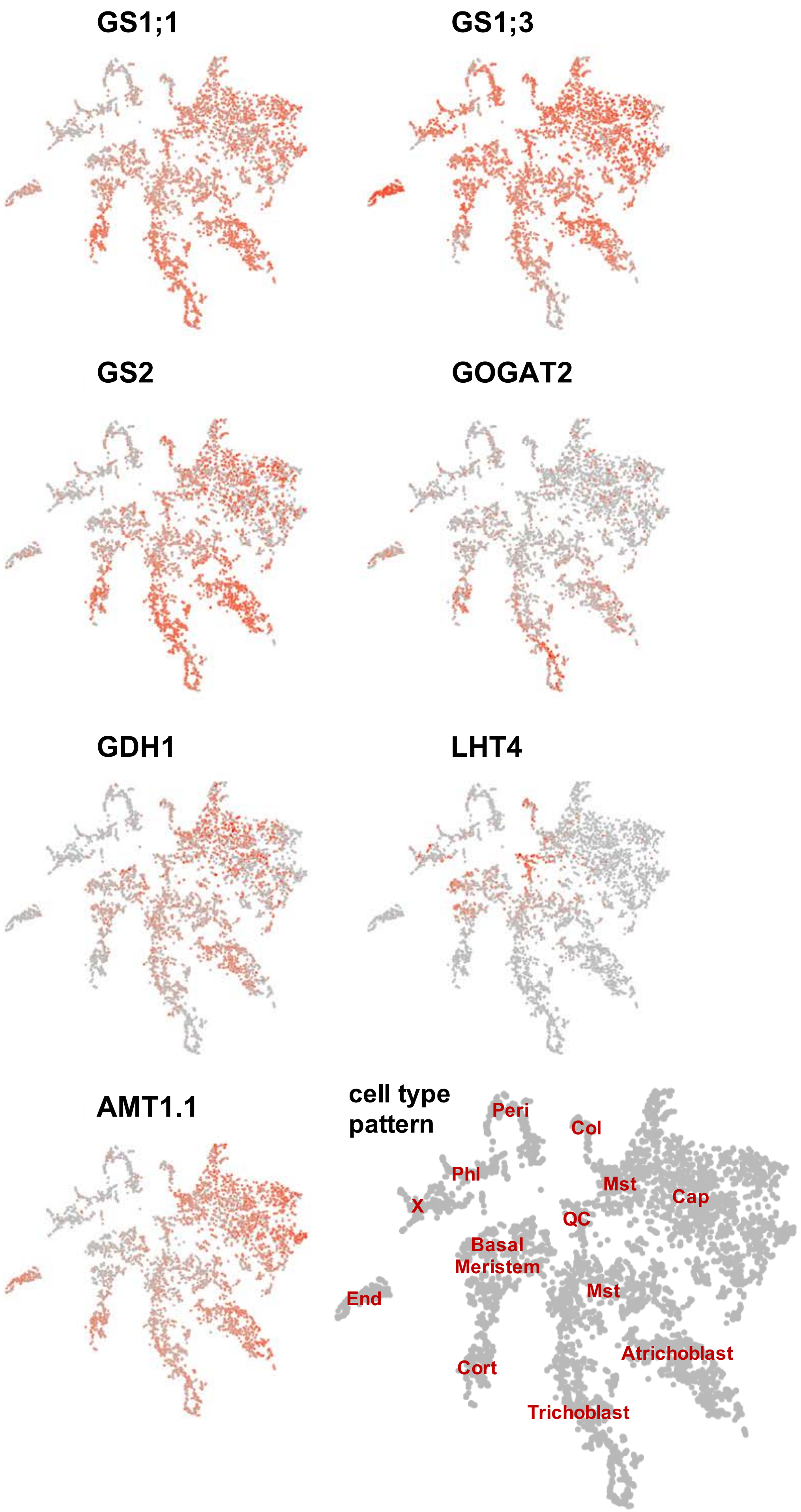
GS1;1
GS1;3
GS2
GOGAT2
GDH1
LHT4
AMT1.1
cell type pattern
Peri
Col
Phl
Mst
Cap
X
QC
Basal Meristem
End
Mst
Cort
Atrichoblast
Trichoblast

**Figure S11. t-distributed Stochastic Neighbor Embedding (t-SNE) representation of single cell transcriptional profiling from Arabidopsis roots.** Single cell RNA-Seq (scRNA-Seq) analysis reveals cell and developmental stage-specific mRNA profiles for select glutamate-associated genes: *GLUTAMINE SYNTHETASE* (*GS*) isoforms *GS1;1* (At5g37600), *GS1;3* (At3g17820) and *GS2* (At5g35630), *GLUTAMATE SYNTHASE 2* (*GLUTAMINE-α-OXOGLUTARATE AMINOTRANSFERASE 2*, *GOGAT2*, At2g41220), *GLUTAMATE DEHYDROGENASE 1* (*GDH1*, At5g18170), putative *LYSINE/HISTIDINE TRANSPORTER 4* (*LHT4*, At1g47670), and the ammonium transporter *AMT1.1* (At4g13510) in roots. Red letters indicate cell identity of select regions; QC – Quiescent Center/Niche; Col – Columella; Peri – Pericycle; Phl – Phloem; - X – Xylem; End – Endodermis; Cort – Cortex; Cap – Lateral Root Cap. Transcripts for *GOGAT1* were not detected by scRNA-Seq, while transcripts of *GLUTAMATE RECEPTOR-LIKE* genes were lowly abundant and lacked specific expression patterns (not shown).
